# Beyond Pairwise Interactions: Charting Higher-Order Models of Brain Function

**DOI:** 10.1101/2025.06.24.661306

**Authors:** Andrea Santoro, Matteo Neri, Simone Poetto, Davide Orsenigo, Matteo Diano, Marilyn Gatica, Giovanni Petri

## Abstract

Traditional models of brain connectivity have primarily focused on pairwise interactions, overlooking the rich dynamics that emerge from simultaneous interactions among multiple brain regions. Although a plethora of higher-order interaction (HOI) metrics have been proposed, a systematic evaluation of their comparative properties and utility is missing. Here, we present the first large-scale analysis of information-theoretic and topological HOI metrics, applied to both resting-state and task fMRI data from 100 unrelated subjects of the Human Connectome Project. We identify a clear taxonomy of HOI metrics — redundant, synergistic, and topological—, with the latter acting as bridges along the redundancy–synergy continuum. Despite methodological differences, all HOI metrics align with the brain’s overarching unimodal-to-transmodal functional hierarchy. However, certain metrics show specific associations with the neurotransmitter receptor architecture. HOI metrics outperform traditional pairwise models in brain fingerprinting and perform comparably in task decoding, underscoring their value for characterizing individual functional profiles. Finally, multivariate analysis reveals that — among all HOI metrics — topological descriptors are key to linking brain function with behavioral variability, positioning them as valuable tools for linking neural architecture and cognitive function. Overall, our findings establish HOIs as a powerful framework for capturing the brain’s multidimensional dynamics, providing a conceptual map to guide their application across cognitive and clinical neuroscience.

---

The brain’s remarkable computational and adaptive abilities have been traditionally explored using network models [1–5], where nodes represent distinct brain areas and connections denote pairwise relationships between them. Structural connectivity (SC) maps the anatomical wiring of the brain [6, 7], while functional connectivity (FC) captures statistical dependencies in neural activity [8–10]. Over the past two decades, these network models have significantly advanced our understanding of brain function in health [11–13], during tasks or cognitive states [14–19], and in clinical populations [20–23]. However, by focusing solely on dyadic interactions, they often miss the complex group-level dynamics underpinning processes like rapid network reconfiguration during decision-making and learning, or the evolution of brain dynamics across age [24–27].

Mounting evidence from micro-to macro-scale studies [28–31] increasingly highlights the importance of higher-order interactions (HOIs) [32–34] — the collective dynamics involving simultaneous interactions among three or more brain regions [35–40]. Such interactions underpin crucial neural mechanisms, including neuronal avalanches, cell assemblies, and distributed encoding [10, 25, 27, 39, 41–43]. Recent advances in information theory and algebraic topological approaches provide robust theoretical and computational frameworks for characterizing higher-order properties, thereby bridging traditional network models and the emergent spatiotemporal complexity of neural systems [44–48].

Building on seminal works [49, 50], information-theoretic approaches provide a robust framework for capturing the collective dynamics of neural systems, by characterizing the interactions among brain regions in terms of different kinds of information: *redundancy* and *synergy* [51–53]. Redundant interactions reflect duplicated or overlapping information between regions, often within the same functional module — such as sensorymotor systems — and are hypothesized to support robust and fault-tolerant coding [27, 46, 54]. In contrast, synergistic interactions arise when information is accessible only through the joint activity of multiple regions, revealing computations not captured by individual regions [37, 44]. These typically span distinct networks, including the default mode and frontoparietal systems, and have been linked to cognitive integration, flexibility, and consciousness [46, 55, 56]. While it is known that conventional FC analyses predominantly capture redundancy [43, 54, 57–59], synergy is increasingly recognized as critical for distributed computation [25, 60– 63], development [64], aging [24, 65, 66], neuromodulation [67, 68], and it is clinically relevant in conditions such as Alzheimer’s disease [69, 70], stroke [71], schizophrenia [72], and autism [73].

Parallel to these developments, topological data analysis (TDA) [74–76] has emerged as a powerful tool to probe the geometrical and topological structure of highdimensional neural data [77, 78]. By leveraging concepts from algebraic topology [79] — such as persistent homology [80, 81] and Reeb-like graphs [82, 83] — TDA detects invariant features, like homological cycles, which reveal hidden higher-order relationships within brain networks [84–91]. A growing body of literature shows that these approaches capture aspects of the brain’s architecture that are invisible by traditional pairwise analyses, offering fresh insights into emergent properties relevant to consciousness [85, 92], health [93], cognition [47] and disease [94–96].

However, the growing interest in HOIs and the resulting rapid development of diverse methodological frame-works — especially in information theory and topological data analysis — has led to a fragmented research landscape. Existing approaches often operate in isolation, each grounded in distinct theoretical assumptions and computational principles. As a result, it remains unclear how these methods relate, whether they capture overlapping or complementary aspects of brain organization, and under what conditions one may be preferred over another.

Here, we address this critical gap by providing the first systematic comparison of higher-order frameworks in the human brain. We evaluated ten widely used metrics — spanning information-theoretic and topological approaches — and benchmarked them against classical pairwise connectivity using resting-state and task-based fMRI data from 100 unrelated subjects in the Human Connectome Project [97, 98]. By mapping their relationships, spatial profiles, and behavioral relevance, our analyses uncover three distinct clusters of HOI metrics and reveal how they align with known cortical hierarchies and micro-architecture. While pairwise functional connectivity performs comparably in task decoding, higher-order metrics provide added value in brain fingerprinting and brain–behavior mapping. Together, this work provides a first insightful lens to navigate the growing landscape of higher-order analyses and establishes a foundation for their broader application in cognitive and clinical neuroscience.

## RESULTS

To provide a comprehensive benchmark of higher-order descriptors of human brain organization, we analyzed resting-state and task fMRI data from 100 healthy unrelated subjects of the Human Connectome Project (HCP) [97, 98]. Following standard preprocessing, we extracted regional time series using a combined parcellation that included 100 cortical regions from the Schaefer atlas [99] and 16 subcortical regions from the Tian atlas [100], yielding 116 regional signals per subject (see Methods). This whole-brain representation serves as the foundation for computing a range of higher-order metrics spanning from information theory to topological data analysis.

As a baseline measure of pairwise interdependence, we consider functional connectivity (FC) as the Pearson correlation between all pairs of regional time series [101, 102]. This widely used measure captures zero-lag linear relationships and serves as a reference point for evaluating more complex, higher-order descriptors.

Next, we quantified redundant and synergistic dependencies using three complementary information-theoretic frameworks, among the most employed in the field. First, we used the Integrated Information Decomposition (PhiID) in its bivariate form [46, 103], estimating synergy (*PhiID Syn*) and redundancy (*PhiID Red*) as information that persists from past to future between pairs of regions. Second, we employed Partial Entropy Decomposition (PED) [57, 104] on triplets of regions to isolate the synergistic (*PED Syn*) and redundant (*PED Red*) components of their joint entropy. Third, we calculated the O-information [50], a multivariate measure quantifying the net balance between synergy and redundancy in triplet interactions; negative values (*O-Syn*) indicate synergy-dominated relationships, while positive values (*O-Red*) reflect redundancy dominance. Each redundant-synergistic pair is hereafter referred to as a conjugate metric, reflecting complementary dimensions of a single decomposition framework.

In parallel, we examined topological features of brain interactions. From the FC matrix, we derived two homological scaffolds [85]: the frequency-based scaffold (*Fscaff*), which weights edges by the number of onedimensional loops they appear in, and the persistence-based scaffold (*Pscaff*), which weights edges by the persistence of these loops. Both approaches identify edges contributing to persistent homological features, i.e. one-dimensional loops, thereby retaining the most topologically informative connections [85]. Finally, to capture higher-order temporal structure, we relied on a recent topological approach [47, 105] to identify hypercoherent triangles (*Triangles*) — triplets of mutually coherent interactions irreducible to pairwise links — and tracked the evolving homological scaffold over time (*TScaffold*). Together, these metrics — spanning topological and information-theoretic approaches — provide a comprehensive lens to characterizing higher-order interdependencies in large-scale brain organization (Table I; Fig. 1). Although applications here focus on triplets for consistency and tractability, most frameworks are extensible to larger sets of regions (See Methods for a detailed description of each framework).

**Table I.** Overview of the metrics to characterize large-scale brain organization. Each method is classified by its analytical domain (Type: Classic, Topology, or Information Theory [IT]) and whether it incorporates temporal information (Temporal). While all genuine higher-order metrics were applied to triplets for consistency, most are extensible to larger region sets. Time-resolved metrics were averaged across time to enable comparison with static descriptors (see Methods).

| Method | Type | Temporal |
| --- | --- | --- |
| FC [1] | Linear | ✗ |
| Pscaff, Fscaff [85] | Topology | ✗ |
| *PhiID Red, PhiID Syn* [103] | IT | ✓ |
| *O-Red, O-Syn* [50] | IT | ✗ |
| *PED Red, PED Syn* [57] | IT | ✗ |
| TScaffold, Triangles [47, 105] | Topology | ✓ |

**Figure 1.**
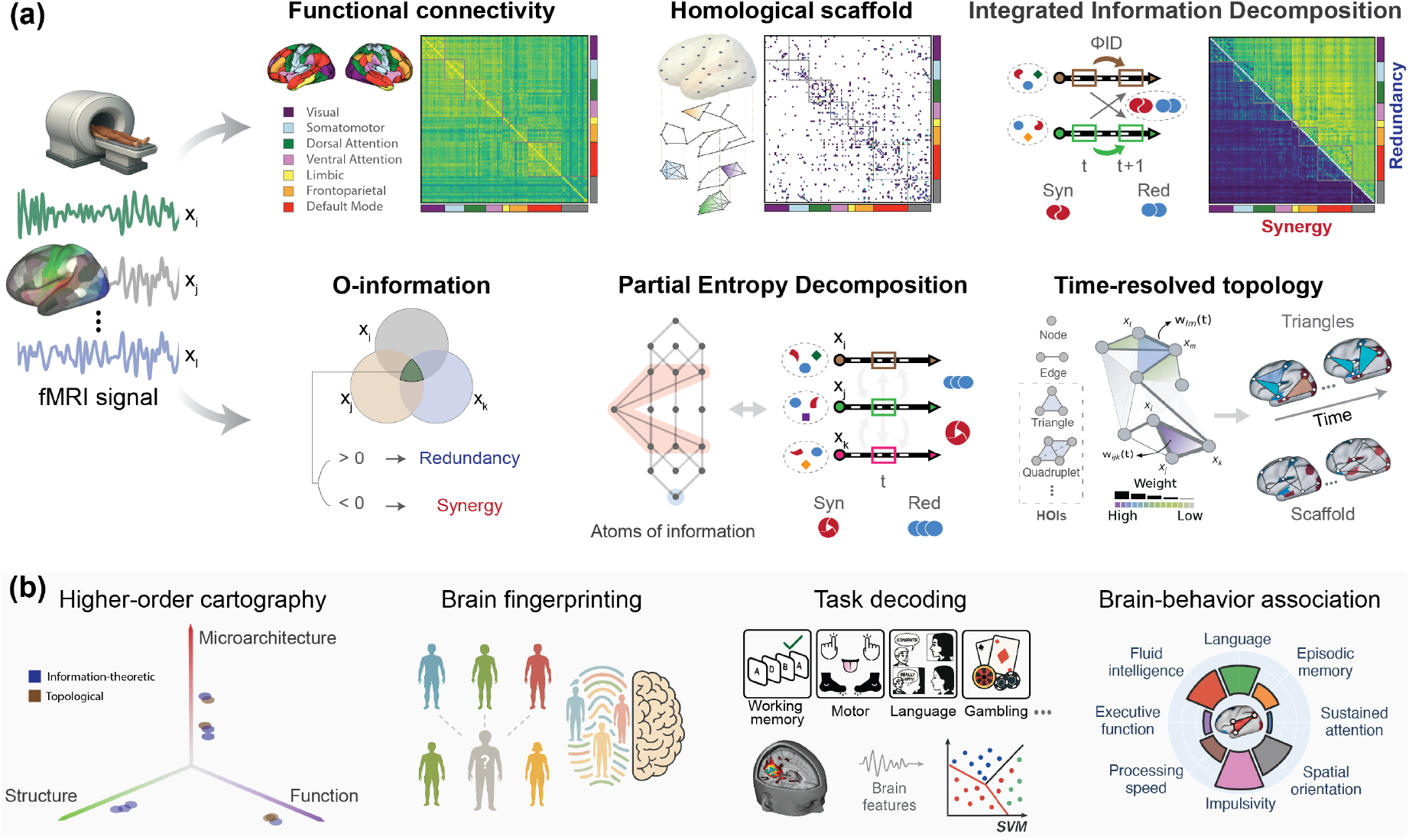
Overview of the approaches and applications. **(a)** Schematic illustration of the metrics used to characterize functional relationship from fMRI data. Approaches include functional connectivity (FC), homological scaffolds derived from persistent topology (Pscaff, Fscaff), and the bivariate form of Integrated Information Decomposition (PhiID), which partitions information into redundant and synergistic components. Other approaches explicitly capture three-way dependencies, including O-Information, Partial Entropy Decomposition (PED), and a time-resolved topological method that tracks hypercoherent triangles (Triangles) and the evolving scaffold over time (TScaffold). **(b)** We consider several brain applications, including the structural–functional cartography of higher-order organization, subject-level brain fingerprinting, cognitive task decoding, and associations between brain organization and behavioral traits. Information-theoretic and topological descriptors offer complementary perspectives on brain micro-architecture, function, and individual variability.

### Higher-order taxonomy

We begin by examining how each connectivity metric maps onto the brain’s large-scale organization and how they relate to one another in terms of spatial distribution and informational content (Fig. 2). To enable consistent comparisons across methods, we projected all metrics onto brain regions and pairwise connectivity by aggregating weights into region-wise nodal scores and edge-wise connectivity scores, respectively (see Methods for details). Figure 2**a** shows the resulting connectivity matrix and cortical maps, averaged across resting-state fMRI data of the 100 subjects (each with 2 sessions and 2 phase-encoding directions, see Methods for details). All values are expressed as percentile ranks to enable comparison across methods.

**Figure 2.**
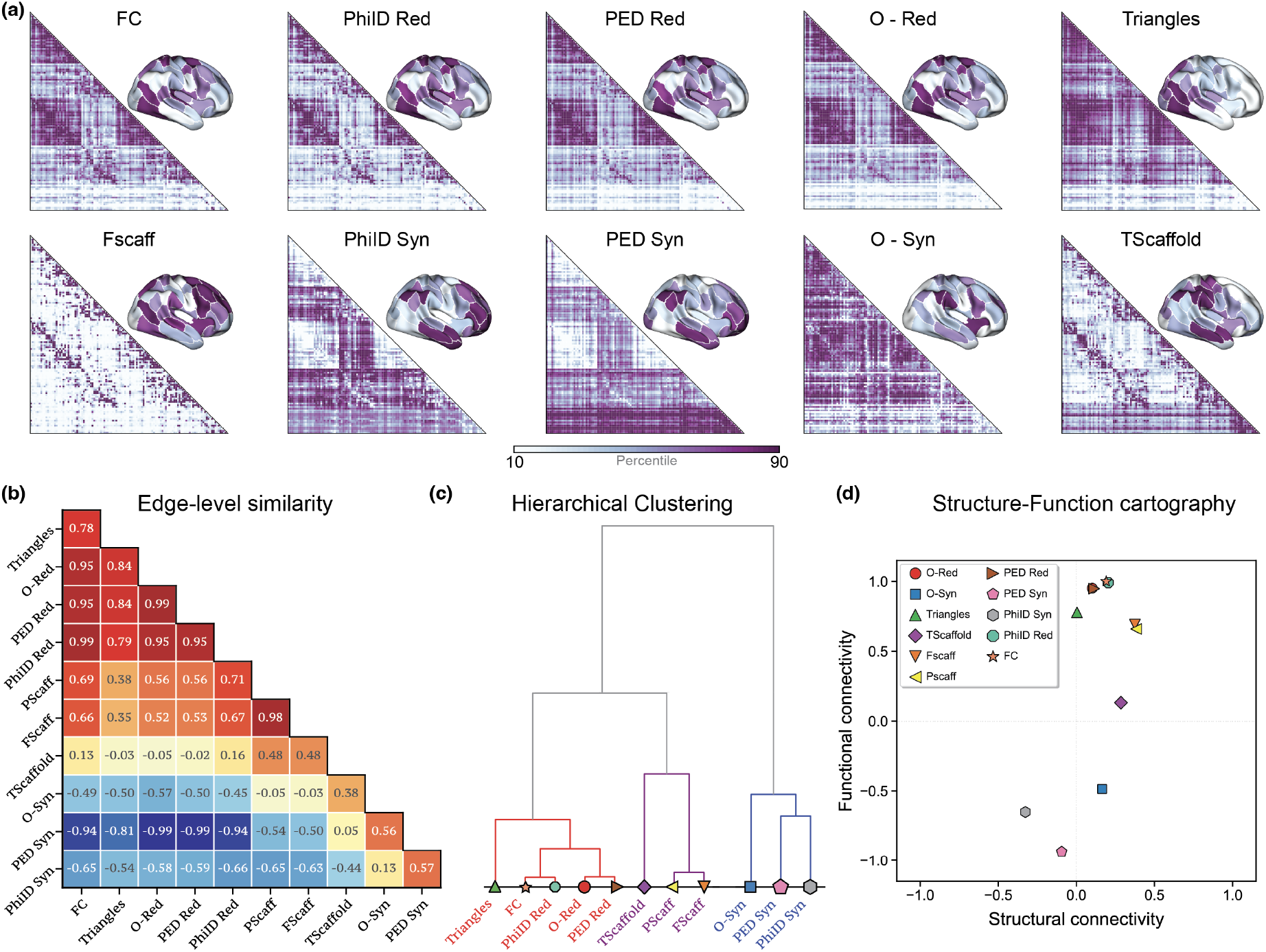
Navigating the landscape of higher-order brain observables. **(a)** Nodal- and edge-level spatial distributions of the connectivity metrics examined in this study, averaged across 400 resting-state fMRI scans (100 subjects, each with 2 sessions and 2 phase-encoding directions, see Methods for details). Metrics shown in the first row predominantly highlight contributions from sensory-motor regions, whereas those in the second row exhibit more distributed patterns, both in nodal values and edge-level connectivity. **(b)** Edge-level similarity between all metrics, computed using Spearman correlation, reveals clusters of methods with shared spatial structure. **(c)** A Ward-linkage hierarchical clustering based on a distance metric 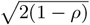 shows a clear separation between redundancy, topological, and synergistic descriptors, confirming the complementarity of their underlying representations. **(d)** Structure–function cartography positions each measure in relation to group-average structural and functional connectivity, highlighting differences in their alignment with anatomical and functional architecture, revealing distinct groupings.

The spatial patterns reveal distinct organizational signatures. As shown in the top row of Fig. 2**a**, metrics such as FC, redundancy-based observables (e.g., PhiID Red, PED Red, O-Red), and triangle-based metrics tend to highlight sensory and visual cortices and connections between regions belonging to the same resting state network (RSN) [106]. This is shown in SI Fig. S1 (nodal projections by RSN) and SI Fig. S2 (within vs. between-RSN connectivity). In contrast, metrics that capture synergistic interactions (e.g., PhiID Syn, PED Syn) can be associated with interactions between regions belonging to different resting state networks and assign greater weight to transmodal systems, including the frontoparietal and default mode networks. These findings align with prior observations linking redundancy to stable, stimulus-driven processing in unimodal systems, and synergy to integrative processing in transmodal regions involved in abstract cognition [46, 57]. Yet, this is the first time these principles are systematically confirmed across a unified set of classical, topological, and information-theoretic metrics.

To quantitatively assess the relationships between the different metrics while preserving the richness of the original data, we computed pairwise Spearman correlations across their edge-level projections (Fig. 2**b**). The resulting similarity matrix revealed a structured organization, which we visualized through ward-linkage hierarchical clustering of the corresponding distance matrix (Fig. 2**c**). This analysis uncovered three major clusters. The first comprises redundancy-dominated metrics — including FC, PhiID Red, PED Red, O-Red, and Triangles — which capture overlapping or shared information between regions. A second cluster consists of the three metrics based on the topological scaffold (Pscaff, Fscaff, and Tscaffold), reflecting mesoscale structural features. The third cluster groups synergy-dominated metrics — PhiID Syn, PED Syn, and O-Syn — highlighting connections that contribute to unique integrative information. However, this clustering structure shifts when looking at the nodal-level projections: Tscaffold aligns more closely with synergy-based metrics than with topological or redundant metrics (see SI Fig. S3 for the nodal-level counterpart of Fig. 2b-d).

To further investigate how these metrics relate to canonical models of brain organization, we computed their correlations with group-average structural and functional connectivity (Fig. 2**d**). This structure–function cartography places each measure in a two-dimensional space defined by its correspondence with structural connectivity (SC; x-axis) and functional connectivity (FC; y-axis). Redundancy-based metrics —including FC, PhiID Red, PED Red, and Triangles — cluster in a region with strong correlation to FC but weaker links to structure. Pscaff and Fscaff also align with FC, but exhibit stronger coupling to SC, suggesting a link between the topological relations between the functional links and the underlying physical structure of the network. In contrast, synergy-based metrics — including PhiID Syn, PED Syn, and O-Syn — are located in a distinct quadrant characterized by negative correlations with FC and low positive and negative correlations with SC. This decoupling suggests that synergy captures integrative dynamics not accessible with the classical FC approach. Remarkably, the scaffold-based metric (TScaffold) occupies an intermediate position, with a clear shift in the clustering when transitioning from nodal-to edge-level projections, bridging the gap between redundant and synergistic frameworks (see SI Fig. S4 for comparison between nodal and edge projections). This suggests that the Tscaffold might encode a blend of local redundancy and distributed synergy, representing the correlation architecture by which the structural connectivity sustains different kinds of synergy in brain function. Together, these findings reveal that connectivity metrics cluster according to their unique properties, each capturing distinct but complementary dimensions of brain organization.

### Multimodal Characterization of Topological and Informational Gradients

Previous work showed how macroscale cortical gradients derived from conjugate information-theoretic metrics of synergy and redundancy reveal fundamental principles of brain organization [46, 57]. Building on this foundation, we systematically compared informational gradients computed via three complementary frameworks — phiID, PED, and the O-information metric — and a novel topological gradient based on scaffold and triangles metric [47, 105] to explore the relationship between informational and topological gradients. Specifically, for each brain region *i*, we computed the four gradients as the difference between synergy and redundancy in each conjugate metric (e.g., PhiID Syn(*i*) − PhiID Red(*i*)), as well as the difference between TScaffold and Triangles (Fig. 3**a**).

**Figure 3.**
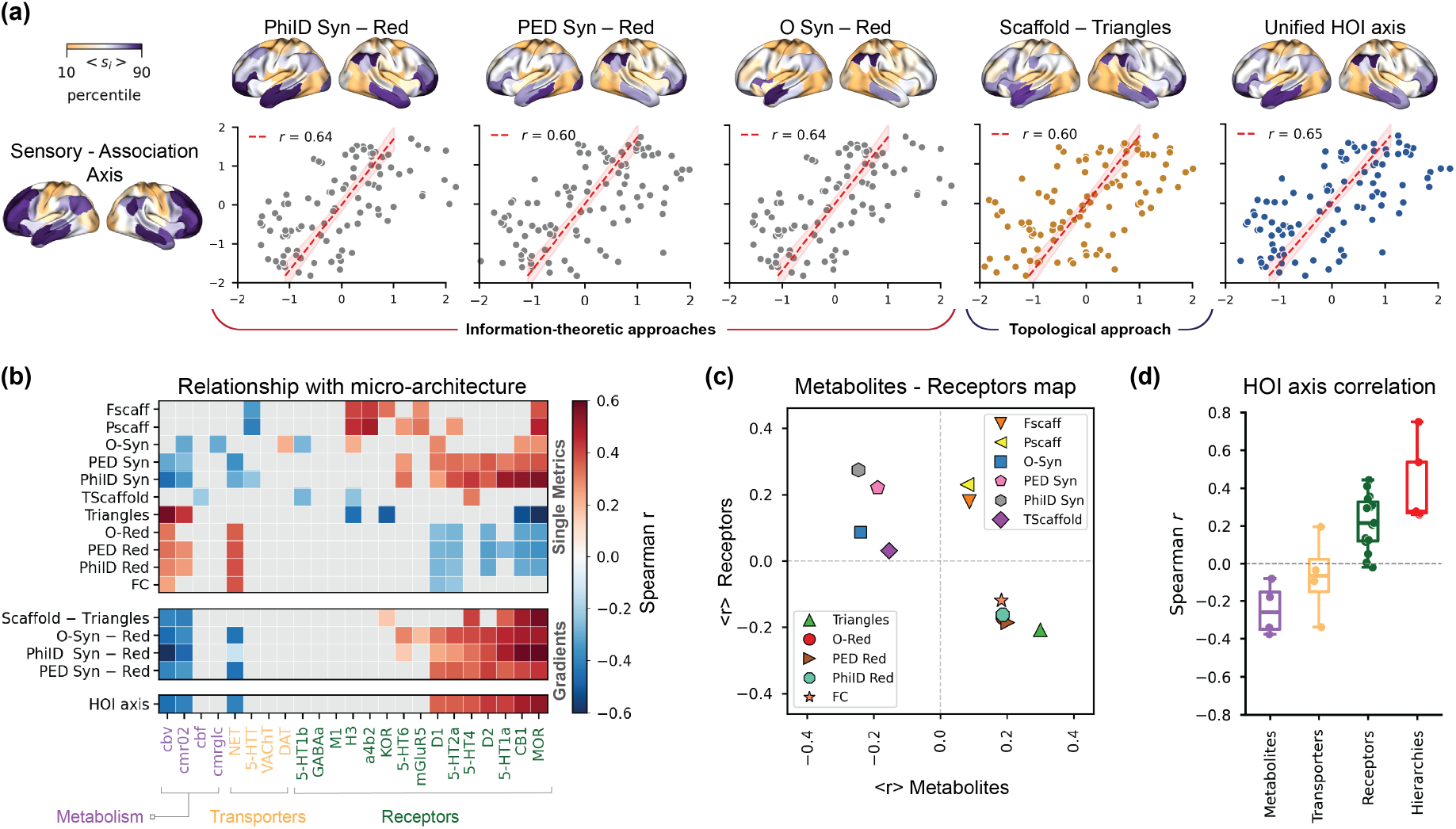
Multimodal characterization of informational and topological gradients. **(a)** Gradients derived from conjugate information-theoretic and topological metrics exhibit a spatial organization that closely aligns with the sensory-association (S-A) axis [107]. The upper panel shows subject-averaged cortical maps for four individual gradients, as well as their mean, which defines the unified higher-order interaction axis (HOI axis). Scatter plots illustrate the relationship between each gradient and the S-A axis, where each brain region is positioned by its R–S value (x-axis) and S-A value (y-axis). **(b)** Correlation profiles between each gradient (y-axis) and cortical maps, including measures of receptors, transporters, and metabolism (x-axis) reveal distinct molecular signatures associated with each gradient. Grey entries indicate non-significant correlations (*p >* 0.05) using spin-permutation test (see Methods) [108]. **(c)** A two-dimensional cartography where each metric is positioned by its average Spearman correlation with receptors (y-axis) and metabolism (x-axis), revealing distinct clusters associated with redundancy, synergy, and topology. **(d)** The HOI axis shows a positive correlation with receptor distributions and cortical hierarchies (see Methods), and a negative correlation with metabolism.

Nodal-level projections of conjugated metric pairs converged on similar spatial patterns. That is, all four gradients peaked in frontoparietal regions and diminished in sensorimotor areas, with strong inter-gradient similarity (Pearson’s *ρ >* 0.8). Furthermore, each gradient aligns strongly with the established sensorimotor–association (S–A) axis, recently defined as the average of multiple cortical hierarchies [107]. All gradients were positively correlated with the S–A axis (Spearman’s *r >* 0.6, *p <* 0.001). On this basis, we can summarize all the gradients above by considering a unified higher-order interactions axis (HOI axis), defined as the mean of these four gradient differences (Fig. 3**a**). This unified gradient provides a single metric that integrates informational and topological signatures of cortical organization (see also SI Fig. S5 for these results at the individual level).

To investigate the micro-architectural properties of these gradients, we correlated each nodal projection with the cortical maps of four metabolic markers, four transporters, and 15 receptors — following recent approaches [109, 110] (Fig. 3**b**). This analysis reveals a notable dichotomy: redundancy- and triangle-based metrics exhibited stronger associations with metabolism, whereas synergy- and scaffold-based metrics aligned more closely with receptor densities . Consistent with these results, the redundancy–synergy gradients and the composite unified HOI gradient correlated positively with receptor maps and negatively with metabolism. Significance was assessed against autocorrelation-preserving spin-based null models [108, 111]

As shown in Fig. 3**c**, the different molecular profiles of our metrics can be captured in a two-dimensional cartography, where each metric is positioned according to its average correlation with metabolism maps (x-axis) and receptor expression (y-axis). This cartography reveals three distinct clusters: (i) a *synergy group* — comprising PhiID Syn, PED Syn, O-Syn, and TScaffold — characterized by strong positive correlations with receptors and negative correlations with metabolism maps; (ii) a *scaffold-specific group* — including scaffold frequency and scaffold persistence (P/Fscaff) — positively correlated with both receptors and metabolism maps; and (iii) a *redundancy group* — consisting of PhiID Red, PED Red, O-Red, and Triangles — aligned positively with metabolism maps and negatively with receptors.

Finally, to contextualize the HOI axis within the broader landscape of cortical organization, we examined its correspondence with four established macroscale hierarchies: evolutionary and developmental cortical expansion [112, 113], T1w/T2w ratio as a proxy for cortical myelination [114, 115], and functional specialization [116]. As shown in Fig. 3**d**, we observed robust positive correlations between the HOI axis and both receptor profiles and cortical hierarchies, while the correlations with metabolic attributes were consistently negative. Thus, this axis not only integrates complementary informational and topological signatures, but crucially it also mirrors the fundamental principles shaping cortical structure. The convergence of these two elements reinforces the view that redundancy and synergy are deeply embedded within the hierarchical organization of the brain.

### Brain fingerprinting and task decoding

We next evaluate how each connectivity measure supports two key applications in functional neuroimaging: *brain fingerprinting* [117], the ability to reliably identify individuals based on their functional connectivity patterns, and *task decoding* [14, 118, 119], the classification of ongoing cognitive states from brain activity.

To evaluate brain fingerprinting performance, we employed two complementary metrics. The identification rate quantifies the ability to correctly match the same subjects across sessions, capturing the proportion of accurately identified individuals within a dataset [117]. In parallel, individual identifiability (*I*_*diff*_) metrics the distinctiveness of a subject’s connectivity profile, defined as the difference between average within-subject and between-subject similarity [120]. These two metrics together provide a comprehensive view of individual trait stability and separability. As shown in Fig. 4**a**, when considering whole-brain connectivity, PhiID Red achieves the highest identification rates, nearing perfect subject matching. Fscaff and FC follow behind. However, when evaluated by *I*_diff_, the highest scores are observed for the synergy and redundancy components of O-information.

**Figure 4.**
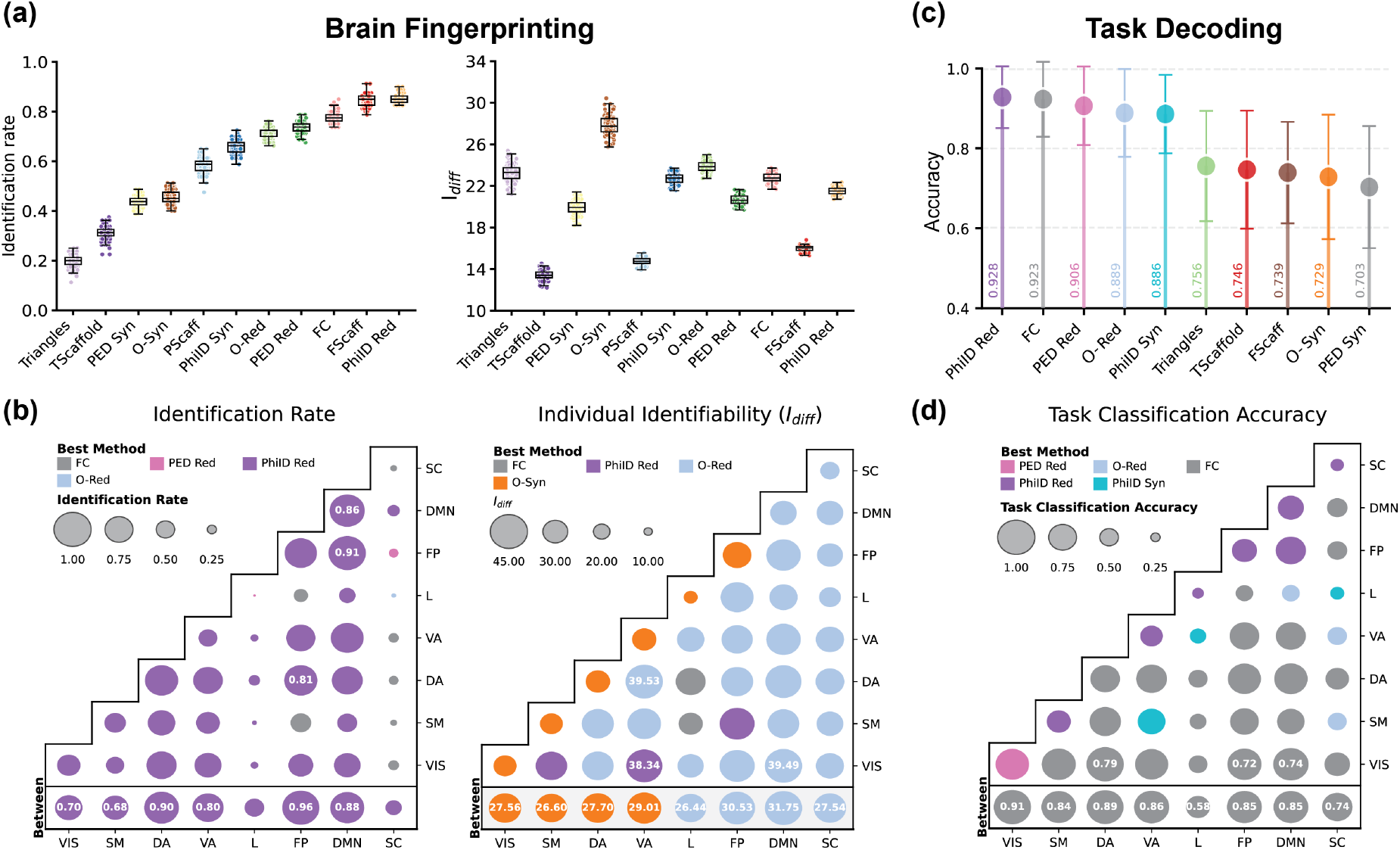
Comparison of connectivity metrics for brain fingerprinting and task decoding. **(a)** Functional brain fingerprinting performance measured via identification rate (left) and individual identifiability score *I*_diff_ (right), using whole-brain connectivity. While PhiID Red and scaffold-based metrics achieve the highest identification rates, redundancy- and synergy-based metrics yield higher *I*_diff_, suggesting they better capture group-level separation. **(b)** Region-level fingerprinting performance, computed separately within and between the seven Yeo resting-state networks [106]. The best-performing method varies by metric and network: functional connectivity (FC) is no longer dominant, with information-theoretic descriptors (e.g., PhiID Red, Red, Syn) showing improved identifiability, especially in transmodal regions. **(c)** Task decoding accuracy using a support vector machine (SVM) in a leave-one-subject-out cross-validation, based on whole-brain connectivity. FC consistently performs among the top methods. **(d)** Task classification accuracy at the resting-state network level, again showing strong performance for FC, particularly in transmodal and associative systems. Together, these results highlight a trade-off: while information-theoretic metrics enhance subject identifiability, FC remains the most effective for decoding task states.

This suggests that O-information metrics better distinguish individuals at the group level, as reflected in the larger separation between intra- and inter-subject similarity. In other words, group-level separability comes at the cost of higher within-subject variability, which undermines identification accuracy.

To examine the spatial specificity of these results, we also assess fingerprinting performance within and between canonical resting-state networks (Fig. 4**b**). While no single metric dominates across all networks and metrics, PhiID Red achieves the best identification rates across most of the systems. In contrast, synergy- and redundancy-based metrics (O-Syn and O-Red) again lead in individual identifiability, particularly within transmodal networks such as the frontoparietal and default mode systems. These findings align with prior evidence that higher-order cognitive areas contribute most strongly to inter-individual variability in brain organization [117, 120–122]. Notably, restricting our analyses to transmodal cortex was almost as discriminative as wholebrain data, reinforcing the view that cognitive individuality is rooted in distributed, high-order networks.

We next assessed the ability of each connectivity measure to decode cognitive states from fMRI data. Following recent approaches [123], we trained a support vector machine (SVM) to classify task conditions using a leave-one-subject-out cross-validation strategy, applied at both whole-brain (Fig. 4**c**) and network-level (Fig. 4**d**) connectivity. Across tasks and brain networks, FC consistently ranks among the best-performing metrics, with redundancy-based approaches (PhiID) leading the pack.

Together, these findings reveal a clear dissociation between connectivity features that support individual identification and those that decode cognitive states. Higher-order metrics — particularly those based on PhiID redundancy, frequency scaffolds, and O-information syn-ergy and redundancy — enhance subject discriminability. In contrast, classical pairwise approaches such as FC remain most effective for decoding dynamic brain states. These results emphasize the importance of aligning connectivity metrics with the specific goals of neuroscientific inference, whether focused on trait-level individuality or state-level cognition.

### FC decomposition and brain-behavior analysis

To comprehensively evaluate how higher-order connectivity metrics contribute to explaining pairwise FC patterns in both resting-state and task fMRI, we defined row-wise multilinear regression models aimed at reconstructing the FC matrix. Specifically, for each brain region *i*, we predict its FC profile — that is, its connectivity to all other regions *j* ≠ *i* — using the different higherorder profiles as predictors.

Regression coefficients were estimated using ordinary least squares, and the goodness-of-fit for each region was quantified by the adjusted *R*^2^, reflecting how well the model predicted that region’s observed FC profile (SI Fig. S6 for a schematic illustration). This regression approach builds on previous methodologies [124– 126], uniquely focusing on individual node-level connectivity. As expected, the model that includes all HOI metrics explained almost fully the FC (range 0.90-0.99). To disentangle the relative contribution of each measure, we performed dominance analysis to partition the total variance explained (*R*^2^) into distinct contributions from three classes of metrics: redundant, synergistic, and topological (Fig. 5**a**).

**Figure 5.**
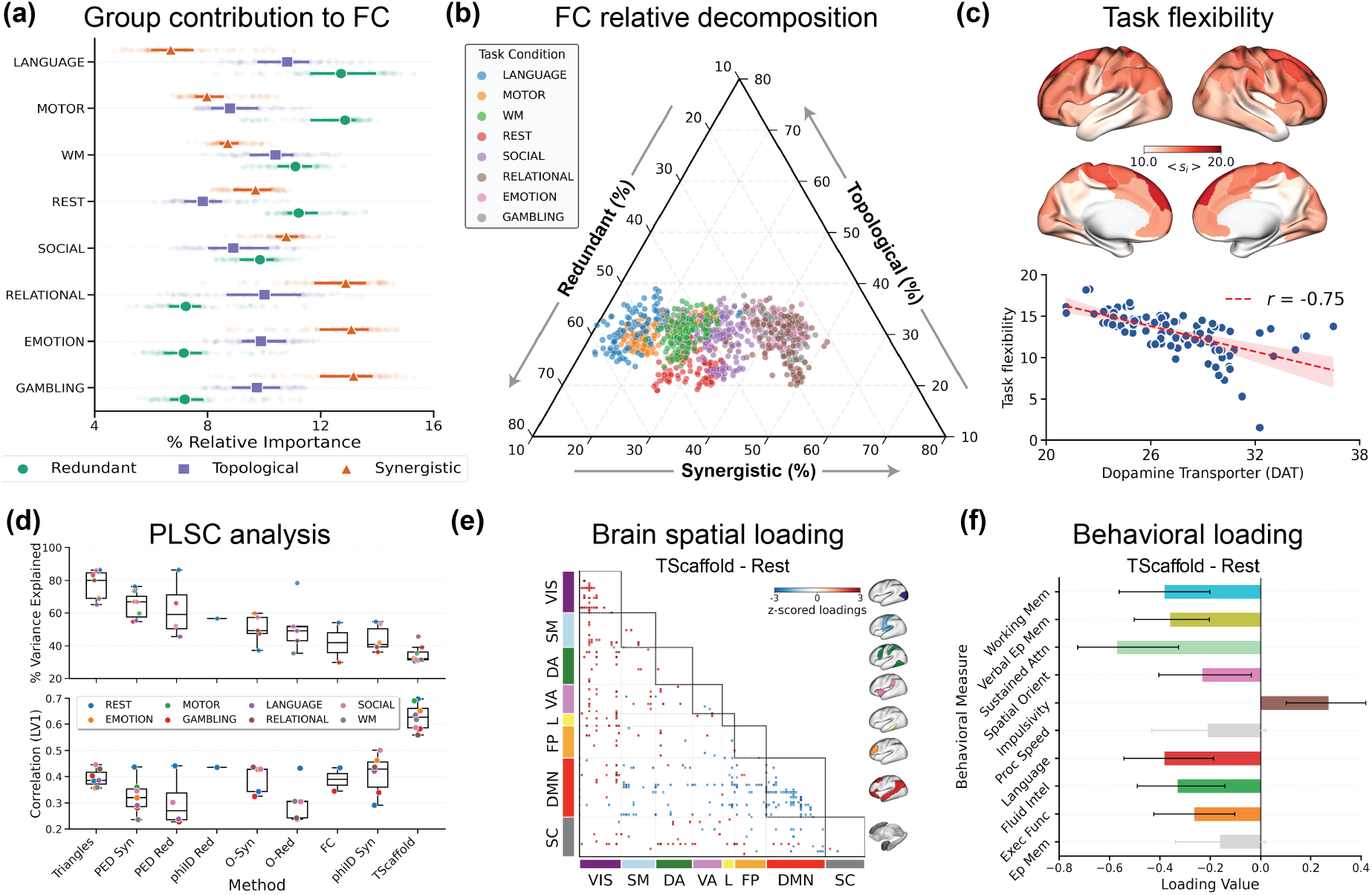
Multivariate analyses reveal task-dependent contributions and behavioral relevance of higher-order metrics. **(a)** Dominance analysis quantifies the relative contribution of three families of higher-order metrics — redundant, synergistic, and topological — in predicting region-wise FC across tasks. While topological metrics contribute consistently, synergistic components dominate in socially relevant tasks (e.g., emotion, gambling), and redundant components are more prominent during rest, motor, and language conditions. **(b)** Ternary plot of brain regions colored by task, showing the normalized contributions of the three metric classes per region and task. Clustering patterns highlight task-specific functional configurations. **(c)** Cortical map of task flexibility, defined as the average distance of a region’s task-specific positions from its centroid in the ternary space (from panel (b)). Regions with higher flexibility show greater task-driven reconfiguration. Flexibility is negatively correlated with dopamine transporter (DAT) density, suggesting a stabilizing role of dopaminergic tone in shaping functional adaptability. **(d)** Variance explained (top) and brain–behavior correlation (bottom) from the first latent variable (LV1) of partial least squares correlation (PLSC), performed independently across tasks and metrics. Topological methods, particularly Triangles and TScaffold, outperform traditional FC in explaining covariance between connectivity and cognitive scores. **(e)** Spatial distribution of the strongest brain loadings for the TScaffold-rest condition, highlighting positive contributions in visual cortices and negative ones in the default mode network (DMN). **(f)** Behavioral loadings associated with LV1 for TScaffold-rest. Negative loadings dominate across multiple cognitive domains, indicating that reduced transmodal connectivity and increased unimodal coupling are linked to lower cognitive performance. Error bars reflect 95% confidence intervals from bootstrapping.

Our analysis revealed distinct patterns across tasks. Topological contributions remained relatively stable across cognitive conditions, whereas redundant and synergistic contributions varied substantially by task type. Specifically, synergistic contributions were most pronounced during “social” cognitive tasks (e.g., emotion, gambling, relational tasks), while redundancy dominated during rest, motor, and language conditions. We visualize these differences using a ternary plot (Fig. 5**b**), where each region’s profile for a given task is positioned based on the normalized contribution of the three metric classes. This visualization qualitatively reveals distinct clusters corresponding to different cognitive tasks. To further explore how each region reconfigures across tasks, we defined a measure of task flexibility: the average distance of a region’s task-specific positions from its centroid in the ternary space. High flexibility indicates greater modulation of a region’s functional profile across cognitive states. Intriguingly, mapping this flexibility across the cortex (Fig. 5**c**) revealed a striking spatial pattern: task flexibility was negatively correlated with dopamine transporter (DAT) maps. In other words, brain regions with lower DAT density exhibited greater shifts in their functional contributions across tasks, suggesting that dopaminergic tone may act to stabilize or constrain higher-order functional reconfiguration. This highlights a potential neurochemical mechanism underpinning regional adaptability in large-scale brain dynamics. Finally, to determine whether certain methods better capture brain–behavior relationships, we performed partial least squares correlation (PLSC) analyses [127, 128]. This multivariate technique identifies latent components that maximize covariance between functional metrics and cognitive performance [129]. We independently applied PLSC to each metric across all tasks, assessing how effectively each method explained individual differences in behavior.

The PLSC analysis consistently identified a single statistically significant latent variable for most methods (Fig. 5**d**), reflecting multivariate associations between functional brain profiles and behavioral outcomes. Topological representations consistently outperformed traditional FC, explaining greater shared covariance and correlations. Specifically, triangles captured the highest covariance (approximately 80% across tasks), followed by PED redundancy and PED synergy (60–70%), while FC explained only 40–50%. Statistical significance was established through permutation testing (1,000 repetitions).

Interestingly, although the triangles approach captured the most covariance, scaffold-based representations yielded the strongest correlations between brain connectivity and cognitive scores. Figure 5**e**–**f** illustrates the loadings of the first significant latent variable for the scaffold-rest condition. This apparent dissociation reflects a key property of PLSC: covariance can be high even when correlation is modest if the latent variables have large variance, whereas smaller latent variances can yield high correlations despite lower shared covariance. In our case, brain regions with the highest positive loadings (top 5%, warm colors) were predominantly located in visual cortices, whereas the highest negative loadings (cool colors) clustered primarily within the default mode network (DMN). Behaviorally, cognitive domains including Working Memory, Verbal Episodic Memory, Sustained Attention, Spatial Orientation, Language, Fluid Intelligence, and Executive Function also showed negative loadings. Concordance in sign indicates a positive brain–behavior association: individuals with reduced scaffold-derived connectivity in transmodal DMN regions and increased connectivity in unimodal visual areas tend to perform worse on a broad set of cognitive tasks. This pattern was not significant for Processing Speed and Episodic Memory (see SI Fig. S7 for analogous results across other methods, where associations were generally weaker).

## DISCUSSION

Recent strides in information theory and computational topology have increasingly offered novel approaches to characterize multiregional higher-order interactions (HOIs) from neuroimaging recordings [27, 44, 130], extending beyond standard pairwise functional connectivity (FC). While individual HOI methods have shown promise in capturing complex neural dependencies overlooked by traditional FC, the field still lacks a systematic understanding of how these metrics relate to one another, how they complement or extend classical connectivity metrics, and what unique neurobiological insights they offer.

Here, we addressed this gap by mapping the landscape of diverse HOI metrics, characterizing their interrelations, biological underpinnings, and evaluating their utility in three key neuroimaging applications: brain fingerprinting, task decoding, and brain–behavior associations. Our comparative analysis, at nodal and edge-level projections, consistently reveals three families of observables: redundancy-based metrics, synergy-based metrics, and topological scaffolds (Fig. 2).

These findings are notable in two major respects. First, despite their distinct theoretical principles, information-theoretic approaches — including O-Information [50], Partial Entropy Decomposition (PED) [57, 104], and Integrated Information Decomposition (PhiID) [103] — converge reliably into synergy and redundancy clusters. This convergence suggests that the brain conveys different kinds of synergy and redundancy with a similar spatial and network organization. While previous work hinted at this alignment [46, 48, 57], here we systematically validated it across multiple HOI frameworks. Second, topological descriptors consistently align with informational modes: triangles align with redundancy, scaffolds correlate with synergy, particularly at the nodal level. This cross-framework correspondence suggests that distinct topological motifs may implement different modes of information processing, reflecting underlying structure–function relationships in the brain.

These observations are consistent with previous studies linking synergy negatively with both FC and structural connectivity (SC) [46], whereas redundancy and triangles positively correlate with these traditional connectivity metrics. Notably, homological scaffolds [47] exhibit a uniquely dual role: nodally, they reflect synergy; at the edge level they correlate strongly with FC and SC. This suggests that scaffolds function as integrative structures, balancing anatomical constraints with dynamic information integration.

Our findings further align with recent topological HOI research: while Varley et al. reported that O-information synergistic interactions are particularly represented in cycles and three-dimensional cavities of data clouds [131], Poetto et al. [132] demonstrated that (significantly larger than expected) PhiID synergistic contributions appear on edges internal to topological cycles. By extending these observations to our definitions of scaffolds, we suggest that the association between topological scaffolds and synergistic interactions reflects a general organizational principle of higher-order neural dynamics.

Yet, what are the biological underpinnings of these organizational common profiles? And, do consistent patterns emerge across different frameworks? To address this, we examined how nodal projections of multiple metrics relate to cortical hierarchies, receptor expression, and metabolism maps (Fig. 3). Remarkably, synergy/scaffold and redundancy/triangle gradients converge onto a shared HOI axis, which strongly aligns with the well-known Sensorimotor–Association (S–A) cortical hierarchy [107]— suggesting that computational modes (synergy vs. redundancy) are embedded within evolutionary, anatomical, and functional gradients [116, 133].

Examining the biological substrates of the HOI axis revealed robust associations with cortical neuro-chemistry. The axis correlated positively with neurotransmitter receptor density, and negatively with metabolic maps and transporter distribution, thus suggesting a clear physiological dissociation. This pattern was mirrored across individual metrics: redundancy- and triangle-based ones mapped onto metabolically efficient sensorimotor regions, supporting stable, low-cost communication; synergy- and scaffold-based ones instead were enriched in receptor-dense association cortices. These findings point to two complementary regimes of brain function respectively—one optimized for reliable signaling [109], and a second for flexible, neuromodulator-driven integration [46]. These results are also consistent with prior work linking transmodal regions to complex cognition, executive control, and consciousness [109, 134], as well as with emerging models in which neuromodulation enables dynamic network reconfiguration in support of adaptive behaviour [69, 135].

When considering neuroscientific applications, higherorder connectivity metrics surpassed pairwise FC in brain fingerprinting [117] (Fig. 4). O-Red and O-Syn achieved the largest identifiability scores (*I*_diff_) [120], whereas PhiID-Red and scaffold-based topological metrics produced the highest identification rates, supporting recent findings [93]. These metrics resolve fine-grained, high-order dependencies that conventional FC misses. Notably, restricting our analyses to transmodal cortices was almost as discriminative as whole-brain data, reinforcing the view that cognitive individuality is rooted in distributed, high-order networks [132].

The pattern reverses for cognitive task decoding: classical FC consistently ranked among the top-performing metrics when considering whole-brain connections or distinct functional networks (Fig. 4). Hence, features that maximize trait-level identification differ from those best suited to capture cognitive states. PhiID redundancy, frequency-domain scaffolds, O-information synergy and redundancy enhance the former –subject discriminability–, whereas classical FC is still optimal for the latter –decoding cognitive states. This aligns with extensive evidence indicating that task-related brain reconfiguration can be effectively captured using classical FC analyses, such as increased integration during cognitively demanding tasks like the N-back [19, 136], or decreased segregation in FC during motor execution tasks [17, 137]. Again, these results show that an appropriate alignment of the chosen metric with the inferential goal — trait versus state in this case — is critical for neuroscientific applications.

Multivariate brain–behaviour analyses further underscore the importance of higher-order topology (Fig. 5). Triangle metrics explain the largest overall variance, yet scaffold metrics show the strongest and most generalizable correlations with behavioural performances across tasks. Because scaffolds sit between nodal synergy profiles and the structural constraints of SC and FC, they appear to integrate redundant and synergistic information into network configurations that directly support behavioral variability.

Finally, we characterized task-dependent FC reconfiguration in terms of HOI contributions. By assigning each region a triplet of indices (redundant, synergistic, topological) per task, we find that cognitive demands modulate the balance of interaction types: socially and relationally demanding tasks amplify synergy/scaffold contributions, whereas sensorimotor and language tasks emphasize redundancy and triangles. These results are both in agreement with previous literature [46, 48] and with the alignment between the HOI and S-A axis. This nuanced modulation of HOI profiles underscores the flexibility of the brain’s high-order architecture in supporting context-dependent processing (Fig. 5)

Our findings should be considered in light of several limitations that might offer directions for future work. First, we focused on a subset of widely used information-theoretic and topological frameworks, selected for their prevalence in the neuroscience literature and widespread use in resting-state fMRI research. Nonetheless, recent extensions of O-information [52, 138–143], as well as other metrics that probe the importance of higher-order structure beyond synergy and redundancy [69, 144, 145], may yield complementary insights. Second, our analyses were conducted on fMRI data from healthy subjects of HCP. Assessing the generalizability of our findings to clinical populations and across different age groups remains an important direction for future research [24, 69], as well as their use to study mesoscale brain organization [146]. Third, the relatively slow sampling of fMRI constrains inference on fast neural dynamics; integrating higher-temporal-resolution modalities such as MEG, EEG, or intracranial recordings would offer important complementary perspectives on the dynamics of HOIs [140]. Fourth, redundancy estimates are known to be sensitive to preprocessing choices; although we used established pipelines [147], we did not systematically explore variation across preprocessing strategies, parcellation schemes, or denoising approaches [148, 149]. A comprehensive assessment of these factors would help clarify their impact on the robustness and generalizability of findings related to higher-order connectivity.

In conclusion, our findings suggest that higher-order interactions reveal latent axes of brain organization that are not accessible via pairwise connectivity. These axes, rooted in distinct computational, structural, and molecular substrates, provide a principled framework to interpret integration, segregation, and their behavioral correlates in the human brain.

## METHODS

### Data sources and preprocessing

#### HCP data

We used fMRI data from the Human Connectome Project (HCP), specifically the U100 unrelated subjects subset from the HCP 900 Subjects Data Release [14, 97]. This cohort includes 100 healthy adult participants (54 females, 46 males; mean age = 29.1 *±* 3.7 years), selected to ensure no family relationships. Per HCP protocol, all participants provided written informed consent approved by the Washington University Institutional Review Board. Ethical approval for analyses on the HCP data was obtained from the Swiss Ethics Committee on research involving humans. All procedures adhered to relevant guidelines and regulations.

MRI data were acquired on a 3T Siemens Prisma scanner using the HCP standard scanning protocol. Structural MRI consisted of a high-resolution T1-weighted 3D MPRAGE sequence (TR = 2400 ms, TE = 2.14 ms, TI = 1000 ms, flip angle = 8°, FOV = 224 *×* 224 mm, voxel size = 0.7 mm isotropic). Diffusion-weighted imaging used spin-echo echo-planar imaging (EPI) with three b-value shells (1000, 2000, and 3000 s/mm^2^, 90 directions per shell plus 6 b=0 volumes; TR = 5520 ms, TE = 89.5 ms, flip angle = 78°, FOV = 208 *×* 180 mm). Resting-state fMRI scans (rfMRI REST1 and rfMRI REST2) were collected over two sessions using gradient-echo EPI (TR = 720 ms, TE = 33.1 ms, flip angle = 52°, FOV = 208 *×* 180 mm, voxel size = 2 mm isotropic), each with two phase-encoding directions: left–right (LR) and right–left (RL).

In addition to resting-state scans, seven task-based fMRI paradigms were acquired across two sessions: gambling (tfMRI GAMBLING), relational (tfMRI RELATIONAL), social (tfMRI SOCIAL), working memory (tfMRI WM), motor (tfMRI MOTOR), language (tfMRI LANGUAGE; including story-listening and arithmetic), and emotion (tfMRI EMOTION). Working memory, gambling, and motor tasks were recorded on the first day, while relational, social, language, and emotion tasks were acquired on the second. For each session, both LR and RL phase-encoding acquisitions were used to compute all the different scores. All analyses used the minimally preprocessed HCP data. Full acquisition and preprocessing details are available in the original HCP publications [14, 97].

#### Brain parcellation

We employed a parcellation of the brain into 116 regions of interest, combining 100 cortical parcels from the Schaefer atlas [99] with 16 subcortical regions from the Tian atlas [100]. This combined cortical–subcortical parcellation was used consistently across structural and functional connectivity analyses.

#### Group-level structural connectivity

A group-representative structural connectome was constructed using diffusion-weighted MRI data from the same 100 unrelated HCP subjects. Diffusion scans were processed with MRtrix3, following a standard pipeline comprising multi-shell, multi-tissue response function estimation, constrained spherical deconvolution, and trac-tography with 2 10^6^ streamlines per subject. Cortical and subcortical segmentations based on the Schaefer (100 regions) and Tian (16 regions) atlases, respectively, were used to parcellate the brain and generate individual connectivity matrices. A group-level structural connectome was then computed by averaging across subjects, preserving edge weights corresponding to streamline counts.

#### Functional Data Preprocessing

We used the minimally preprocessed fMRI data provided by the HCP [150], which includes artifact removal, motion correction, and registration to standard space. Key steps in the HCP functional pipeline included volumetric and grayordinate-based spatial preprocessing, and weak high-pass temporal filtering (cutoff *>* 2000 s).

For resting-state fMRI, we applied additional preprocessing: a Butterworth bandpass filter ([0.008 Hz, 0.08 Hz]) was applied in both forward and reverse directions (python nilearn *NiftiLabelsMasker*).

For task fMRI, the same preprocessing steps were applied, with the exception of bandpass filtering, as the optimal frequency bands for task-based signals remain task-dependent [151]. The resulting region-wise time series were used for all functional connectivity and classification analyses.

#### Brain maps from neuromaps

We used the `neuromaps` toolbox (https://github.com/netneurolab/neuromaps) [152] to obtain a set of brain maps capturing cortical hierarchies together with metabolism, and neurotransmitter receptor and transporter brain topographies. Surface-based maps were transformed to the standard `fsLR32k` cortical surface using linear interpolation and parcellated into 100 cortical regions using the Schaefer-100 atlas in `fsLR32k` space [99]. Volumetric data were retained in MNI152 space and parcellated using the Schaefer-100 volumetric atlas.

Maps depicting cortical hierarchies included the T1w/T2w ratio as a proxy for intracortical myelination [114, 115] and the evolutionary and developmental expansion maps [112, 113]. Also from neuromaps, we recover the Margulies gradients as a proxy for the functional hierarchies, and the Metabolic maps include cerebral blood flow, cerebral blood volume, oxygen metabolism, and glucose metabolism [153, 154]. Neurotransmitter maps encompassed densities of 19 different neurotransmitter receptors and transporters spanning 9 neurotransmitter systems: serotonin (5-HT1a, 5-HT1b, 5-HT2a, 5-HT4, 5-HT6, 5-HTT), dopamine (D1, D2, DAT), norepinephrine (NET), acetylcholine (*α*4*β*2, M1, VAChT), histamine (H3), glutamate (mGluR5), GABA (GABAa/BZ), opioid (MOR), and cannabinoid (CB1) [21, 109, 155, 156]. These maps provide a high-resolution, multi-receptor perspective on the molecular architecture of the human cortex [152].

### Higher-order frameworks

#### *O-Information (*Ω): Red *and* Syn

The *total correlation* [157] TC and the *dual total correlation* [158] DTC, are given by the following formulas:

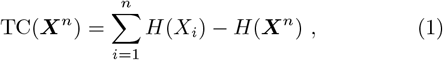

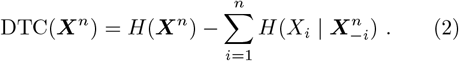

Here, *X*^*n*^ is a vector of *n* random variables, *X*^*n*^ = (*X*_1_, …, *X*_*n*_), *H*(*·*) represents the Shannon entropy, and 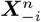 represents the vector of *n* − 1 variables composed by all minus *X*_*i*_ i.e., (*X*_1_, …, *X*_*i*−1_, *X*_*i*+1_, …, *X*_*n*_). Both TC and DTC are non-negative generalizations of mutual information, meaning they are zero if and only if all variables *X*_1_, …, *X*_*n*_ are statistically independent of one another.

For a set of *n* random variables ***X***^*n*^ = (*X*_1_, …, *X*_*n*_), the O-Information [50] (denoted by Ω) is calculated as follows:

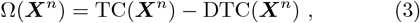

In our study, we employ the O-information to investigate third-order interactions. For three variables *X*_*i*_, *X*_*j*_, and *X*_*k*_, the O-information is equivalent to the interaction information [159] and can be expressed as:

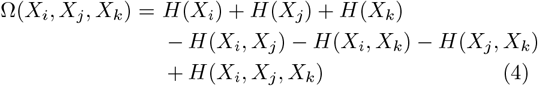

The O-information is a real-valued measure that captures the balance between redundancies and synergies in arbitrary sets of variables, thus extending the properties of the interaction information of three variables [159] to larger sets (see related discussion in Ref. [49]). In particular, the sign of the O-information serves to discriminate between redundant and synergistic groups of random variables: Ω *>* 0 corresponds to redundancy-dominated interdependencies (*O-Red*), and Ω *<* 0 characterizes synergy-dominated variables (*O-Syn*). These metrics were computed using the `HOI` toolbox, specifically the class of functions named `Oinfo` from the repository (https://github.com/brainets/hoi.io) [160].

#### *Integrated information decomposition (Phi ID):* PhiID Red *and* PhiID Syn

Consider three random variables: two source variables *X*^*i*^ and *X*^*j*^, and a target variable *Y* . The *Partial Information Decomposition (PID)* [49] decomposes the total information provided by *X*^*i*^ and *X*^*j*^ about *Y*, given by Shannon’s mutual information *I*(*X\**; *Y*), as follows:

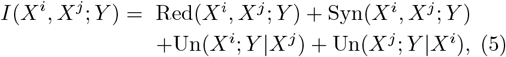

where Red(*X*^*i*^, *X*^*j*^; *Y*) represents the information provided by *X*^*i*^ and *X*^*j*^ about *Y* (redundancy), Syn(*X*^*i*^, *X*^*j*^; *Y*) denotes the information provided jointly by *X*^*i*^ and *X*^*j*^ about *Y* (synergy), Un(*X*^*i*^; *Y* | *X*^*j*^) is the unique information provided by *X*^*i*^ about *Y*, and Un(*X*^*j*^; *Y* | *X*^*i*^) is the information that is provided only by *X*^*j*^ about *Y* . The four terms of this decomposition are naturally structured into a lattice with nodes 𝒜 = {{12}, {1}, { 2}, {1} {2}}, corresponding to the synergistic, unique in source one, unique in source two, and redundant information, respectively. To compute these terms, it is necessary to estimate redundancy. A common approach for this is the Minimum Mutual Information (MMI) estimator, which defines redundancy as the minimum mutual information between each source and the target. Synergy, in contrast, captures the additional information provided by the weaker source when the stronger source is already known [161]. This choice is motivated by the fact that the MMI approach has been benchmarked on Gaussian data [161], and there is a long-standing body of literature supporting the use of Gaussian models to represent fMRI data [162–164].

Consider the stochastic process of two random variables 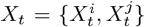 and denote the two variables in a current state *t*, by 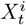 and 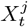, and the same two variables in a past state *t*−*τ*, by 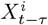^*i*^, and, 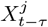. The *Integrated Information Decomposition (*Φ*ID)* is the forward and backward decomposition of 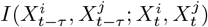, called the time delay mutual information, in redundant, synergistic and unique information [103]. The ΦID can be represented by the forward and backward interactions of the product 𝒜 *×* 𝒜, resulting in 16 distinct atoms: synergy to synergy, redundancy to redundancy, unique in source one to unique in source two (and backward), and redundancy to synergy, among others. Following previous work [46], our analyses focus on two specific atoms quantifying the temporal persistence of redundancy and synergy: the *redundancy to redundancy* atom (*ϕ*_Red_) and *synergy to synergy* atom (*ϕ*_Syn_).

To compute these terms, it is necessary to employ an estimator for the redundancy to redundancy atom (*ϕ*_Red_). Building on the Minimum Mutual Information (MMI) principle and following recent work on this approach, we compute it as follows:

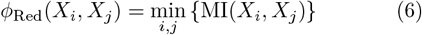

Once this approximation is obtained, all 16 Partial Information Decomposition (PID) atoms can be estimated. In particular, this allows for the computation of the synergy to synergy atom (*ϕ*_Syn_). These metrics were computed using the `HOI` python toolbox, specifically the class `phiID_atoms` from the repository (https://github.com/brainets/hoi.io) [160].

#### *Partial Entropy Decomposition (PED):* PED Red *and* PED Syn

The Partial Entropy Decomposition (PED) framework [57, 104] offers a principled way to decompose the joint entropy of a set of discrete random variables into “atoms” that capture distinct modes of information sharing—specifically, redundancy and synergy. While aligning with the axioms of information decomposition [49, 165]. For a system

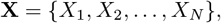

the joint entropy is given by

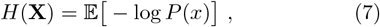

where *x* denotes a particular configuration of **X**.

Central to PED is the idea of “shared entropy”. Rather than summarizing a distribution with a single scalar *h*(*x*) = −log *P* (*x*), this method seeks to determine how different components of *x* jointly exclude probability mass—that is, how they “share” information.

This concept is generalized by considering all non-empty subsets of the system. Given a collection of sources *a*_1_, …, *a*_*k*_ (each *a*_*i*_ representing a subset of components). Redundancy is defined through the measure first proposed by Makkeh et al. [166] and used in Varley et al. [57]:

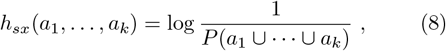

and its expectation yields the global measure

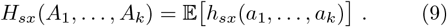

Not every collection of subsets constitutes a valid “atom” in this decomposition. Following [49], Varley et al. [57] restrict their attention to combinations where no source is a subset of any other. Formally, if *s* is the power set of the components (excluding the empty set), the domain of valid atoms is defined as

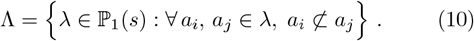

Within the lattice Λ, the partial entropy atom associated with a given collection *α* is obtained via Möbius inversion:

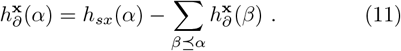

Consequently, the joint entropy can be exactly recovered as

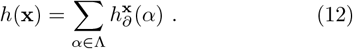

For systems of three variables, PED allows us to decompose classical information-theoretic quantities into contributions attributable to redundancy and synergy. In particular, one may define a measure of *redundant structure* as

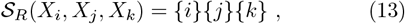

which captures the information redundantly available from each individual variable. In contrast, the *synergistic structure* is given by the sum of all atoms representing information that emerges only when variables are considered jointly:

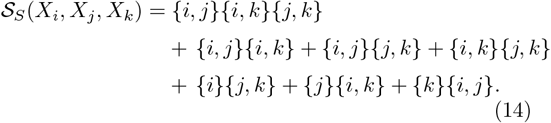

#### Homological Scaffolds: Fscaff and Pscaff

Persistent and frequency-based scaffolds are built on top of FC matrices and designed to highlight mesoscale structures [85]. These methods identify groups of interacting regions by selecting topologically relevant groups of links that collectively form topological loops. The construction of these scaffolds relies on persistent homology, a technique in computational topology that analyzes the shape and structure of high-dimensional datasets [167, 168].

Our approach begins with a weighted functional network, *W* = (*V, E, ω*), where *V* represents vertices (brain regions), *E* represents edges (functional connections), and *ω* : *E →* ℝ denotes the edge weights (i.e. Pearson correlations). To capture the network’s topological features across different intensity scales, we employ a weight rank filtration [169]. This involves constructing a sequence of binary graphs, *G*_*w*_ = (*V, E*_*w*_), where an edge *e ∈ E* is included in *G*_*ω*_ if its weight 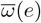 is larger than or equal to a threshold *ω*. For each *G*_*ω*_, we associate its clique complex *K*_*ω*_. A clique complex is a simplicial complex where every *k*-clique in *G*_*ω*_ is promoted to a (*k*− 1)-simplex [169]. The family of complexes {*K*_*ω*_} then defines a filtration, 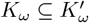, allowing us to track the emergence and disappearance of topological “holes” as the filtration progresses.

We compute the generators 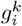 of the *k*-th homological group *H*_*k*_ along this filtration. Each generator identifies a specific hole in the network’s mesoscopic connectivity, such as a one-dimensional cycle (loop). The importance of each hole is quantified by its persistence (*π*_*g*_ = *δ*_*g*_ − *β*_*g*_), where *β*_*g*_ and *δ*_*g*_ are the “birth” (appearance) and “death” (disappearance) weights of the hole during the filtration, respectively.

These persistent homology generators are then projected onto a weighted network to form the homological scaffolds [85]. We defined two types of scaffolds:

- **Frequency-filtered scaffold (*Fscaff*)**: This scaffold assigns weights to connections based on the number of topological loops they are part of. If an edge *e* belongs to multiple generators *g*_*i*_, its weight 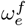 is defined as the sum of indicator functions for each generator it belongs to:

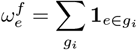

where 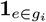 is 1 if edge *e* is part of generator *g*_*i*_, and 0 otherwise. Edges frequently participating in such loops are considered more significant.
- **Persistence-based scaffold (*Fscaff*)**: Here, connection weights are determined by the persistence of the topological loops during the filtration process. If an edge *e* belongs to multiple generators *g*_*i*_, its weight 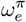 is defined as the sum of the persistence values of those generators:

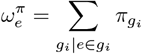

Links contributing to loops that persist over a broader range of filtration values (i.e., have higher 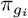) are given higher importance, indicating more robust topological features.

By definition, both scaffolds share the same edge set of links, but with different weighting schemes. This construction highlights the role of links that are part of many and/or long-persistence cycles, effectively isolating functionally important edges within the network.

#### Triangles and temporal scaffold

To characterize higher-order functional interactions over time, we rely on the recent topological approach [105], which generalizes edge-centric analyses [170] to dynamic simplicial complexes based on co-fluctuations among triplets of brain regions. Let 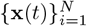 be a multivariate time series with *N* brain regions and *T* time points, where each signal **x**_*i*_ is first *z*-scored to ensure comparability across regions: 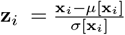. While second-order co-fluctation are defined as a temporal unrapping of the Pearson correlation coefficient [170], i.e. *w*_*ij*_ = *z*_*i*_*z*_*j*_, we define the instantaneous third-order co-fluctuation among regions (*i, j, k*) at time *t* as:

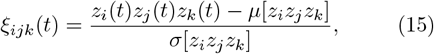

where *µ*[*·*] and *σ*[*·*] denote the temporal mean and standard deviation of the product time series. To differentiate concordant group interactions from discordant ones in a 3-order product, concordant signs are always positively mapped, while discordant signs are negatively mapped. Formally,

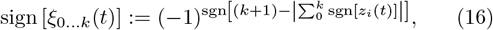

where sgn[•] is the signum function of a real number. As a result, each co-fluctuation is assigned a signed magnitude:

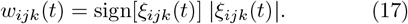

For each time point *t*, the full set of pairwise and triplet co-fluctuations defines a weighted simplicial complex 𝒦^*t*^, consisting of edges (*i, j*) with weight *w*_*ij*_(*t*) = *z*_*i*_(*t*)*z*_*j*_(*t*), and triangles (*i, j, k*) with weight *w*_*ijk*_(*t*) as defined above. In this setting, triangles capture coordinated fluctuations among three regions beyond pairwise correlations.

To extract the mesoscale organization of these timevarying simplicial complexes, we construct a topological filtration, considering multiple thresholds based on the weights of 𝒦^*t*^, generating a sequence of nested simplicial complexes that approximates with increasing precision the original weighted simplicial complex. At each step, triangles violating the simplicial closure condition—i.e., triangles whose minimum edge weight is lower than the triangle weight—are excluded and recorded in a violation list Δ_*v*_ (*Triangles*).

In a similar fashion as described in the previous section, by applying persistent homology to the well-defined filtration constructed from 𝒦^*t*^, we identify generators of one-dimensional cycles and construct a temporal scaffold graph (*Tscaffold*). Each edge weight equals the number of 1D-cycles in which that edge participates, thus highlighting links that frequently close topological loops. We compute both triangle-based and scaffold-based metrics at every time point and derive their static counterparts by averaging these metrics across time. These metrics were computed using the original Julia code from the repository (https://github.com/andresantoro/RHOSTS).

### Nodal and Edge projections

To analyze the information conveyed by the different higher-order metrics at the edge and node levels, we applied a downward projection approach. Specifically, for all metrics defined at the triplets level — including O-information redundancy and synergy, PED synergy and redundancy, as well as the violating triangles — we computed edge- and node-level summaries by aggregating contributions from associated triplets. That is, for each edge (*i, j*) we assign a weight *w*_*ij*_ equal to the average sum of the weights of triplets including that edge, i.e. triplets of the form (*i, j*, •) with a weight *w*_*ij*•_, and the average is computed over the number of triplets *n*_*ij*•_ which include that edge. Formally:

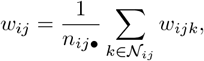

where 𝒩_*ij*_ denotes the set of nodes forming a triangle with edge (*i, j*). Similarly, we define the nodal strength *w*_*i*_ of node *i* as the average sum of weights of the triangles connected to node *i*. Formally:

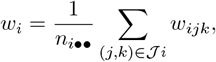

where 𝒥_*i*_ is the set of node pairs (*j, k*) that form a triplet with node *i*. This procedure enables comparison across metrics at a common resolution.

### Clustering

To investigate the similarity structure among the different metrics, we performed hierarchical clustering based on pairwise dissimilarity. We first computed a dissimilarity matrix between all pairs of the 11 edge or nodal projections using the distance metric 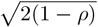 [171, 172], where *ρ* is the Spearman correlation coefficient between each pair of metrics. This measure captures the degree to which two metrics share a similar spatial distribution across functional connections (Figure 2**b**) or cortical regions (SI Figure S3a). We then applied agglomerative hierarchical clustering using the `dendrogram` function from the `scipy.cluster.hierarchy` module in Python. The linkage method used was *ward* linkage, which merges clusters based on minimizing the total within-cluster variance, choosing at each step the pair of clusters whose merger results in the smallest increase in the overall sum of squared distances. The resulting dendrogram provides a data-driven representation of how the metrics group together based on their spatial dissimilarity, revealing shared patterns or functional relationships across different metrics.

### Statistical analysis

To assess the significance of the spatial associations between the nodal projections of our metrics and various cortical maps (including metabolic attributes, receptor densities, and cell-type gene expression), we employed both spatially informed permutation testing and subject-level analyses. Specifically, we used a spin permutation test to evaluate whether observed correlations exceeded what would be expected by chance given the spatial autocorrelation of cortical anatomy. This method preserves the spatial structure of cortical maps while randomizing their alignment through rotations on the spherical surface, generating a null distribution against which empirical correlations can be compared. We performed 1000 spin permutations using the `spin_test` function from the ENIGMA Toolbox (https://github.com/MICA-MNI/ENIGMA) [173], following established procedures for spatial statistical inference in surface-based neuroimaging data. In parallel, we computed Spearman correlations at the individual subject level, correlating each participant’s nodal projection with each cortical map. This produced a distribution of correlation values across subjects, which we tested against zero using two-sided one-sample *t* -tests. To correct for multiple comparisons across brain features, we applied Bonferroni correction. This dual approach allowed us to assess both the spatial robustness and individual-level consistency of our results.

### Partial Least Square Correlation (PLSC)

To examine the relationship between brain activity and behavior across the four different methods, we rely on Partial Least Square Correlation (PLSC) analyses. These analyses aimed to find connections between the functional links of each method and 10 cognitive scores across subjects. For each method, we consider the same number of functional connections, corresponding to 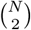 entries.

To enhance the stability of the functional connections, we averaged them over the two resting-state fMRI recordings (i.e., *REST*_1 and *REST*_2). The cognitive scores considered belong to 10 cognitive subdomains in the HCP dataset, covering behavioral traits like episodic memory, executive functions, fluid intelligence, language, processing speed, self-regulation/impulsivity, spatial orientation, sustained visual attention, verbal episodic memory, and working memory [14]. When multiple raw scores were available for a subdomain, we obtained a single score through principal component analysis.

We performed the PLSC analysis separately for each of the methods. PLSC identifies linear combinations of functional connectivity values that maximally covary with linear combinations of cognitive scores, determined through singular value decomposition of the data covariance matrix [128]. The weights of these linear combinations are commonly referred to as brain function and cognitive saliences, representing the left and right singular vectors of the data covariance matrix. We assessed the statistical significance of the PLSC components using permutation testing (1000 permutations; correlation patterns with *p <* 0.05 were considered significant) [128]. To evaluate the reliability of non-zero salience values, we implemented a bootstrapping procedure (1000 random data resampling with replacement) and calculated standard scores with respect to the bootstrap distributions (salience values were considered reliable if the 95% CI did not cross the zero) [128]. We quantified the amount of cognitive traits’ variance explained by functional connectivity values by summing the squared singular values corresponding to the significant PLSC components. This sum was normalized by the total sum of squared singular values obtained for each PLSC analysis [128]. Notice that we rely on the pypls package for computing the PLSC analyses (https://github.com/netneurolab/pypyls).

## Code availability

One of the key outcomes of this work is the development of a collection of ready-to-use code for analyzing time series data from various higherorder interaction perspectives. All code is publicly available in the following GitHub repository: https://github.com/nplresearch/HOI lenses analysis.

The analyses were conducted using Python (tested on version 3.10) for computing the Partial Entropy Decomposition (PED), PhiID, and O-information, and Julia (tested on version 1.10.4) for analyses involving Triangles, Fscaffold, and Pscaff. The implementations of O-information and PhiID are based on the open-source Python toolbox HOI [160], available at https://github.com/brainets/hoi.

## Acknowledgments

M.N. has received funding from the French government through the ‘France 2030’ investment plan, managed by the French National Research Agency (Agence Nationale de la Recherche; references: ANR-16-CONV000X / ANR-17-EURE-0029), as well as from the Excellence Initiative of Aix-Marseille University—A*MIDEX (AMX-19-IET-004). G.P. acknowledges support by ERC Consolidator Grant RUNES (Grant no. 101171380). A.S. has received funding from the European Union’s Horizon 2020 research and innovation programme under the Marie Sklodowska-Curie grant agreement no. 101208090.

## Contributions

A.S., M.G., and G.P. conceived and developed the study idea. A.S. (lead) and M.N. (supporting) performed the formal analysis with contributions from S.P, and D.O.; M.D. preprocessed the fMRI data. A.S., M.N., S.P., D.O., M.G., and G.P. contributed to the interpretation. A.S. and M.N. wrote the original draft with valuable revisions by G.P., M.G., D. O., and S.P. All authors contributed to review and editing. A.S. and M.N. contributed equally to this work as co-first authors; S.P. and D.O. contributed equally.

