## Supplementary Information for "Beyond Pairwise Interactions: Charting Higher-Order Models of Brain Function"

June 24, 2025

### Nodal projection and inter-subject variability

To complement the results presented in the main text, Supplementary Figure 1 shows the distribution of average nodal projection ranks across subjects for each of the seven resting-state networks (RSNs) defined by the functional parcellation of Schaefer et al. [3], along with 16 Subcortical areas (SC) of the Tian atlas [5]. These RSNs include: Visual (VIS), Somatomotor (SM), Dorsal Attention (DA), Ventral Attention (VA), Limbic (L), Frontoparietal (FP), and Default Mode Network (DMN). For each subject, nodal projection values were ranked across all regions from lowest to highest. The ranks of regions belonging to the same RSN were then averaged, yielding a mean rank per RSN per subject. Finally, the across-subject distribution of these average ranks was plotted for each RSN. This analysis was repeated for all the metrics and the results reveal a consistent pattern: unimodal sensorimotor networks tend to rank higher for redundancy-based metrics, whereas multimodal associative networks tend to rank higher for synergy-based metrics. Moreover, we observed that the subject variability is not significantly different between the different metrics.

### Network differences between higher-order descriptors

To further understand how functional interactions distribute across the brain at the edge level, we examined the relative strength of connections occurring within versus between RSNs. This comparison sheds light on whether certain metrics preferentially capture cohesive modules or more distributed connectivity patterns. As illustrated in Fig. S2, redundancy-based metrics and scaffold descriptors exhibit significantly stronger within-network connections, consistent with known modular organization of the brain FC [1]. Conversely, synergy-based metrics, a part O-Syn, exhibit stronger connections between regions belonging to different RSNs, capturing patterns of information integration across distinct functional modules of the brain (Fscaff: $t = 17.50$, $p = 2.56 \times 10^{-59}$; Pscaff: $t = 16.17$, $p = 7.10 \times 10^{-52}$; PED Syn: $t = -17.61$, $p = 1.21 \times 10^{-60}$; PhiID Syn: $t = -24.13$, $p = 9.11 \times 10^{-101}$; TScaffold: $t = 31.89$, $p = 1.95 \times 10^{-153}$; Triangles: $t = 10.77$, $p = 1.05 \times 10^{-25}$; O-Red: $t = 17.59$, $p = 7.56 \times 10^{-61}$; PED Red: $t = 17.78$,

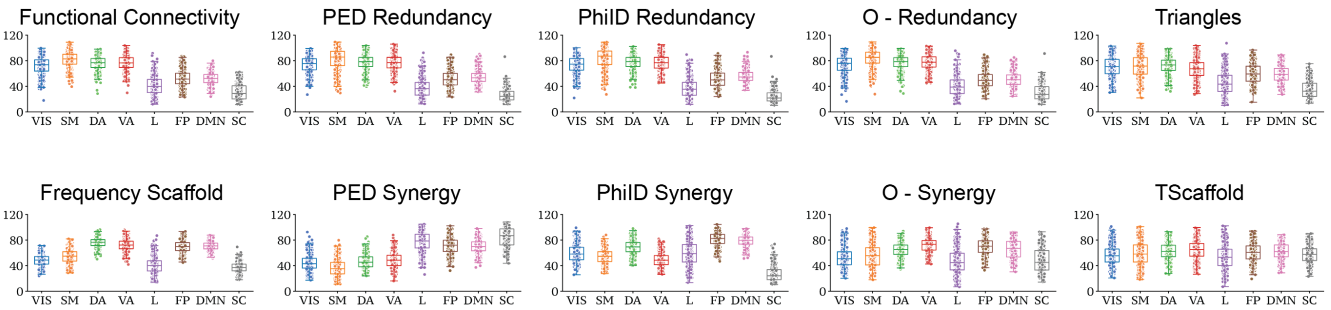


Figure S1: **Comparison of nodal projection across different resting state networks.** This panel displays the distribution across subjects of the average nodal strength for each resting-state network (RSN) and subcortical areas — Visual (VIS), Somatomotor (SM), Dorsal Attention (DA), Ventral Attention (VA), Limbic (L), Frontoparietal (FP), Default Mode Network (DMN) and subcortical areas (SC) — for all metrics under investigation. As *Pscaff* and *Fscaff* yield highly similar results, only *Fscaff* is presented for clarity. Notably, nodal projections reveal opposing trends for redundancies and synergies: synergies exhibit stronger nodal strength in multimodal brain regions, whereas redundancies dominate in unimodal, sensorimotor areas.

$p = 1.38 \times 10^{-61}$; PhiID Red: $t = 25.16$, $p = 1.68 \times 10^{-107}$; FC: $t = 25.10$, $p = 3.87 \times 10^{-109}$; O-Syn: $t = 1.87$, $p = 0.062$). These results highlight a fundamental distinction in the network structure related to different metrics. Notably, our results are consistent with previous observations for PhiID redundancy and synergy [2], while also extending these trends to a wider set of higher-order descriptors.

### Shared spatial structures among metrics

One of the central aims of this study is to determine whether different metrics give rise to similar spatial patterns across the brain. While the main text focuses on edge-level projections—given their capacity to retain more fine-grained information—here we turn to nodal projections to investigate inter-metric similarity at the regional level. We computed pairwise Spearman correlations between nodal projections and applied hierarchical clustering to reveal groups of metrics with similar spatial distributions. As shown in Fig. S3, three well-defined clusters emerge, corresponding to redundancy-based (FC, O-Red, PhiID Red, PED Red, Triangles), topological (Pscaff and Fscaff), and synergy-based descriptors (O-Syn, PhiID Syn, PED Syn, Tscaffold). Notably, Tscaffold, which clusters with Pscaff and Fscaff at the edge level, aligns more closely with synergy-based metrics at the nodal level. This shift underscores how the level of analysis (edge vs. node) can significantly affect metric interpretation and the relationships among descriptors.

### Divergence between edge and nodal projections

To directly examine how metrics diverge across projection levels, we compared their similarity properties at the edge- and node-level. Specifically, we computed the difference between the correlation matrices obtained at each level, capturing how inter-metric relationships shift depending on the projection. In parallel, we assessed the stability of each metric's alignment with FC and SC by comparing correlation values across edge- and node-level projections. Metrics exhibiting minimal change across levels were characterized by similar correlation profiles, while others showed substantial deviation—indicating sensitivity to the level of analysis. Among these, Tscaffold displayed the most pronounced divergence, with marked shifts in both clustering and connectivity correlations. These results reveal that information theoretic metrics and triangles behave quite consistently across levels, whereas the scaffolds, in particular Tscaffold, reflect fundamentally different network properties when evaluated at the edge versus node level projection (Fig. S4). These findings emphasize that edge- and node-level projections offer complementary

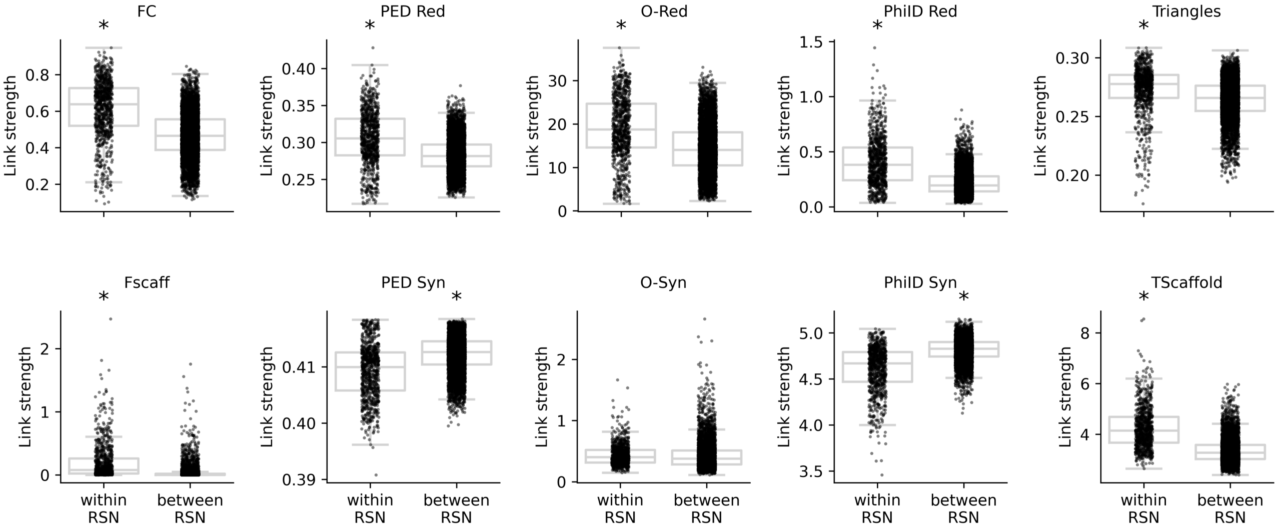


Figure S2: **Comparison of Connection Strengths Within and Between Resting-State Networks.** This panel shows the distribution of link strengths connecting pairs of brain regions within the same resting-state network (RSN) versus those connecting regions across different RSNs. Results are shown for all the metrics under study, since *Pscaff* and *Fscaff* produce highly similar results, only *Fscaff* is shown for visualization purposes. Statistical comparisons between each pair of distributions were performed, consistently yielding $p$-values $< 0.001$ for all the metrics a part O-Syn. All metrics showed a positive $t$-statistic—indicating stronger within-network connections—except for the synergistic metrics.

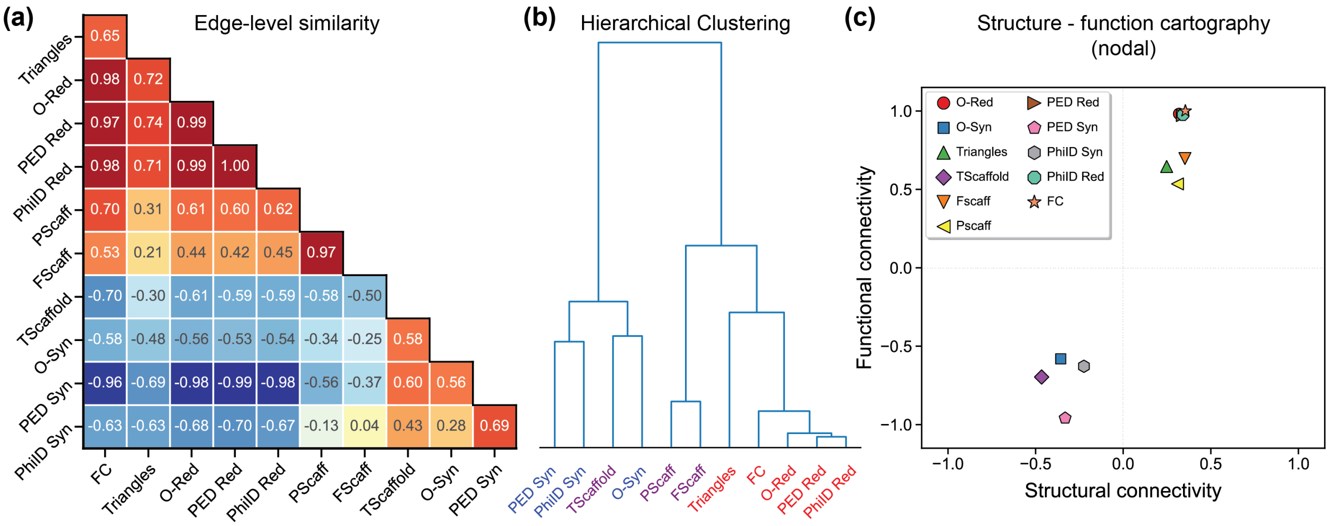


Figure S3: **Clusters at the nodal level.** Similarity between the nodal projection of all measures, computed using Spearman correlation, reveals clusters of methods with shared spatial structure. (a-b) A complete-linkage hierarchical clustering based on a distance metric $\sqrt{1-2\rho}$ shows a clear separation between redundancy, topological, and synergistic descriptors, confirming the complementarity of their underlying representations. Notably this clustering differs with respect to the one computed from the edge level projections (Fig. 2c). Here, Tscaffold belongs to the cluster of synergistic metrics, note that we keep the same color for metrics labels as the one used in Fig. 2. (c) Structure–function as in Fig. 2d but computed at the nodal level. At the nodal level, there are only two clusters, one of synergistic and temporal scaffold nodal projections (anticorrelating with both structural connectivity strength (SC strength) and functional connectivity strength (FC strength), and the other of redundancies and static scaffold correlating with both SC and FC strength.

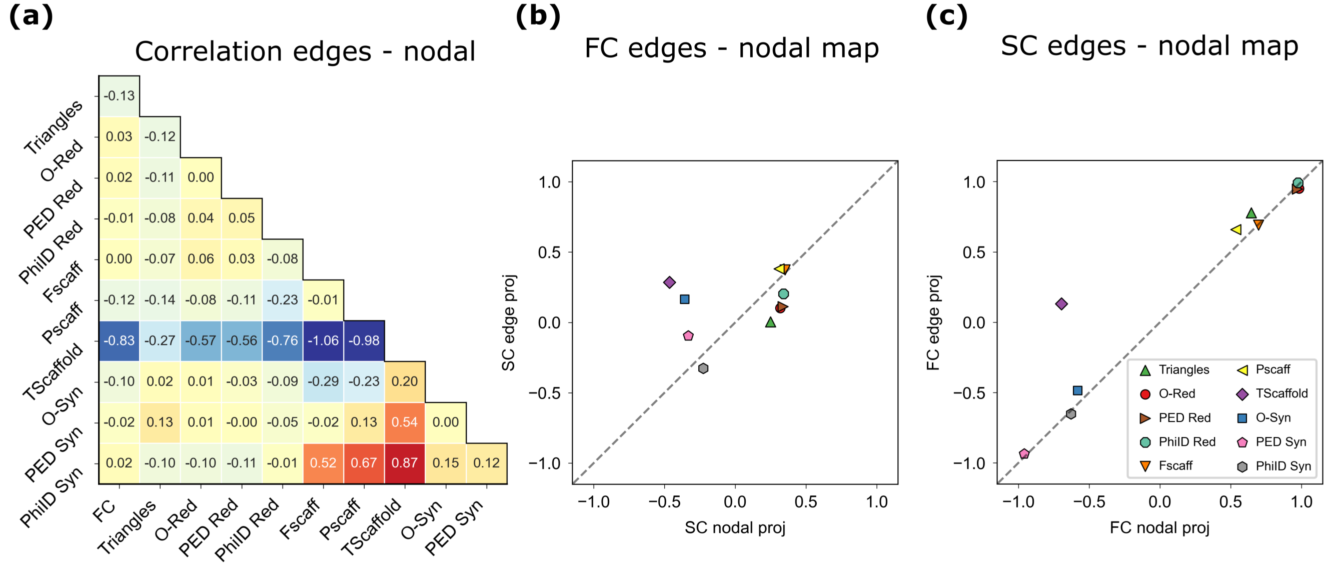


Figure S4: **Comparison of Edge- and Node-level Analyses.** Panel (a) shows the point-wise difference between the correlation matrices of the metrics computed from the edge-level projection and those computed from the node-level projection. This matrix highlights that Tscaffold is the metric whose behavior changes most notably, as it shifts cluster when transitioning from edge- to node-level analysis. In panels (b) and (c), the $y$-axis represents the correlation with FC (panel b) and SC (panel c) computed at the edge level, while the $x$-axis shows the correlation with FC (panel b) and SC (panel c) at the nodal level. The greater a metric's distance from the bisectrix, the greater the difference in its behavior between the two projection levels. Panels (b) and (c) are consistent with what observed in panel (a), confirming the scaffold as the metrics with the strongest variability across projection levels.

perspectives on brain network organization. Furthermore, they highlight the unique role of Tscaffold as a bridge between redundancy and synergy: while it shares an edge-level correlation structure with other topological descriptors, its nodal projection aligns more closely with synergy-based brain topographies. This dual behavior suggests that Tscaffold may serve as an integrative descriptor, capturing relevant correlation structure at the edge level, while involving similar topography as synergy.

### Inter-subjects variability

As described in the main text, to further investigate the relationships between different methods at the nodal level, we employed a range of brain topographies capturing distinct organizational hierarchies, as well as the spatial distribution of various receptors, neurotransmitters, and metabolites. Prior work has shown that gradients derived from conjugate metrics—such as redundancy and synergy—relate to known functional hierarchies. We systematically extended this analysis by incorporating a wider array of redundancy- and synergy-based descriptors. In particular, we focused on the relationship between the nodal projection of the metrics and the sensorimotor–association (S-A) axis proposed by Sydnor et al. [4], which captures a core gradient of cortical functional organization. To complement the results reported in the main text, we computed this correlation at the subject level to test the inter-subject variability of our results. For each metric, we computed the correlation between its nodal projection and the S-A axis on a subject-by-subject basis, resulting in a distribution of correlation values across subjects. This approach enabled us to quantify inter-individual variability in alignment with this functional gradient. A one-sample $t$-test against zero was used to determine whether the observed correlations were consistently positive or negative at the group level. To control for multiple comparisons, Bonferroni correction was applied across

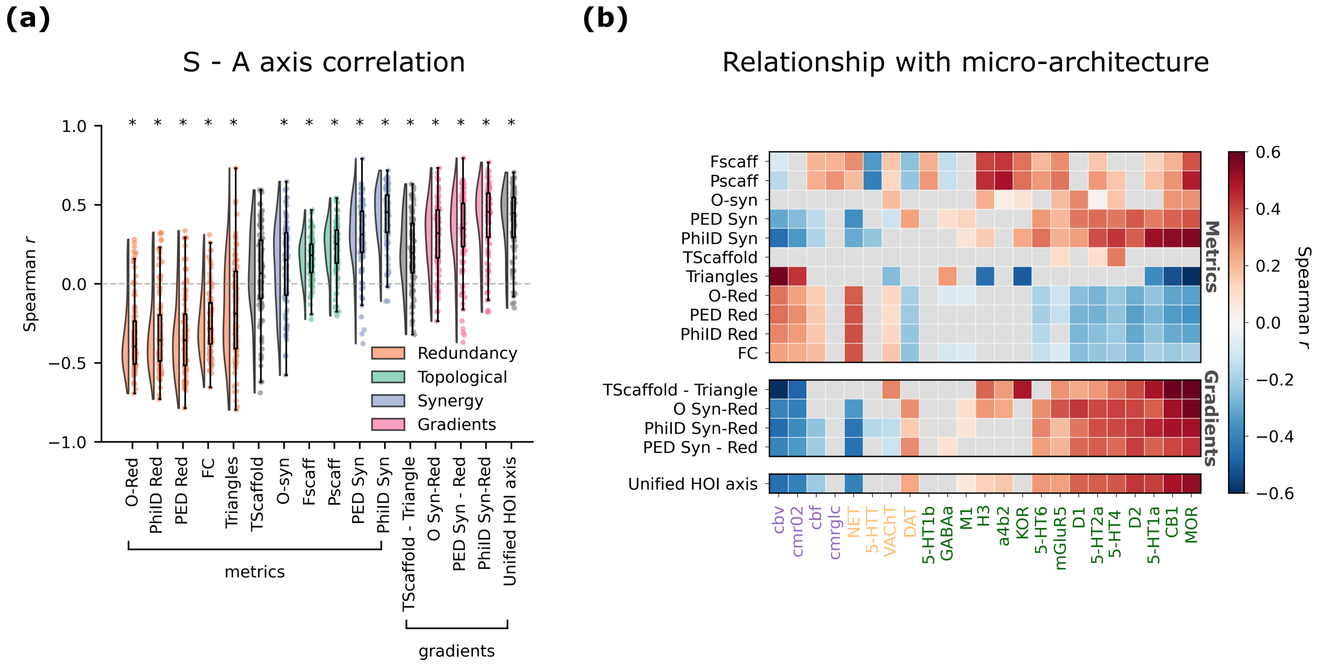


Figure S5: **Subject variability.** (a) Distribution of Spearman correlations between each subject's nodal projection and the Sensorimotor–Association axis defined by Sydnor et al. [4]. Different colors represent different informational or topological metrics. Asterisks denote statistically significant values after Bonferroni correction. (b) Same plot as in Fig. 3b, with the addition that gray squares indicate correlations that were not significant at the population level.

all metrics (Fig. S5a). We applied the same approach to assess how inter-subject variability affects correlations between metric-derived nodal projections and cortical maps of receptor, neurotransmitter, and metabolites density maps. For each map and each metric, subject-level correlations were computed and tested for consistency across the population using a one-sample $t$-test. Results that did not reach significance after Bonferroni correction are shown in grey in Fig. S5b, highlighting variability in how reliably different metrics capture molecular and metabolic spatial gradients across individuals.

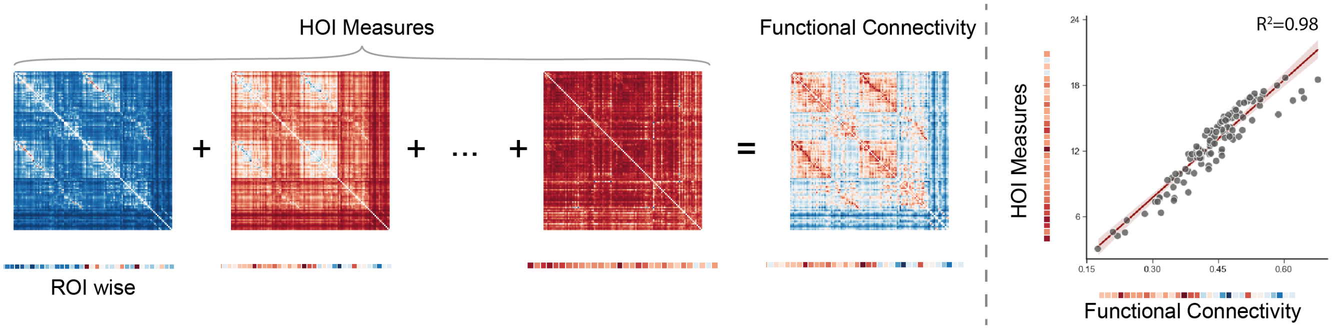


Figure S6: **Node-wise HOI-FC relationship.** Following Vázquez-Rodríguez et al. [6], local structure–function coupling was estimated using a multilinear regression model fit separately for each node $i$. The dependent variable was the functional connectivity $FC_{ij}$ between node i and all other nodes $j \neq i$, and the independent variables were the 10 HOI metrics for each pair $(i, j)$. Model parameters were estimated via ordinary least squares. Goodness of fit for each node was quantified by $R_i^2$ between observed and predicted $FC_{ij}$.

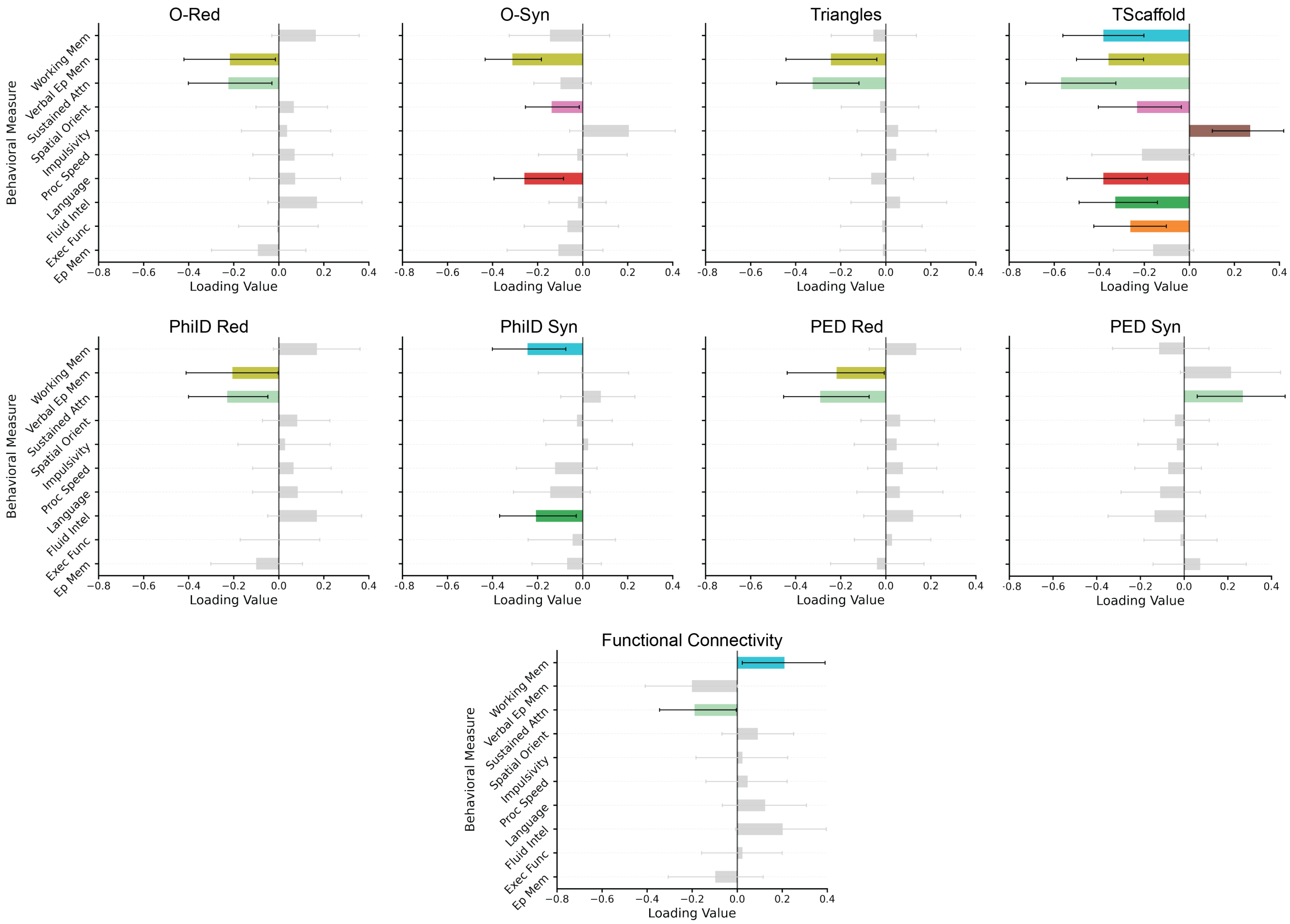


Figure S7: **Behavioral PLSC scores .** Significant behavioral loadings from the Partial Least Squares Correlation (PLSC) analysis linking brain-level metrics to behavioral dimensions. Each panel shows the loading values of behavioral measures for a given brain feature (e.g., O-Red, PhiID Syn, Scaffold, Functional Connectivity), with colored bars indicating statistically significant loadings (bootstrap ratio above 95% confidence interval) and gray bars showing non-significant contributions. Notably, scaffold and synergy-based metrics (e.g., Tscaffold, PhiID Syn, PED Syn, O-Syn) show stronger associations with language and fluid intelligence, while redundancy-based and triangle metrics are more strongly associated with attention and verbal episodic memory.
